# Design of a mucin-selective protease for targeted degradation of cancer-associated mucins

**DOI:** 10.1101/2022.05.20.492748

**Authors:** Kayvon Pedram, D. Judy Shon, Gabrielle S. Tender, Natalia R. Mantuano, Jason J. Northey, Kevin J. Metcalf, Simon P. Wisnovsky, Nicholas M. Riley, Giovanni C. Forcina, Stacy A. Malaker, Angel Kuo, Benson M. George, Caitlyn L. Miller, Kerriann M. Casey, José G. Vilches-Moure, Valerie M. Weaver, Heinz Laübli, Carolyn R. Bertozzi

## Abstract

Targeted protein degradation is an emerging strategy for the elimination of classically undruggable proteins. Here, to expand the landscape of substrates that can be selectively degraded, we designed degraders which are dependent on both peptide sequence and glycosylation status of the target protein. We applied this approach to mucins, O-glycosylated proteins that drive cancer progression through biophysical and immunological mechanisms. Engineering of a bacterial mucin-selective protease yielded a variant for fusion to a cancer antigen-binding nanobody. The resulting conjugate selectively degraded mucins on cancer cells, promoted cell death in culture models of mucin-driven growth and survival, and reduced tumor growth in murine models of breast cancer progression. This work establishes a blueprint for the development of biologics which degrade specific glycoforms of cell surface proteins.

## Introduction

Mucins are glycoproteins that bear a high density of O-glycosylated serine and threonine residues. In species ranging from sea sponges to mammals, mucins are expressed at epithelial and endothelial surfaces, where they defend against physical insults and pathogens^1^. The mechanisms by which mucins exert their functions at these surfaces fall broadly into two categories (Fig. 1a). First, mucins are critical to the initiation and propagation of biophysical signals. For example, their extended and rigid secondary structure enables their use by cells as force-sensitive antennae, as is the case for the mucin MUC1 during integrin-mediated adhesion to soft matrices^2^, and the mucin CD45 during macrophage pinocytosis^3^. Second, the glycopeptide epitopes presented by mucins act as ligands for various receptors, particularly those involved in cell adhesion and immune modulation^4^. For example, the immune cell receptor Siglec-7 (sialic acid-binding immunoglobulin-like lectin 7) binds the mucin CD43 on leukemia cell surfaces and delivers an immune inhibitory signal, analogously to classical checkpoint receptors such as PD-1^5^.

**Figure 1.**
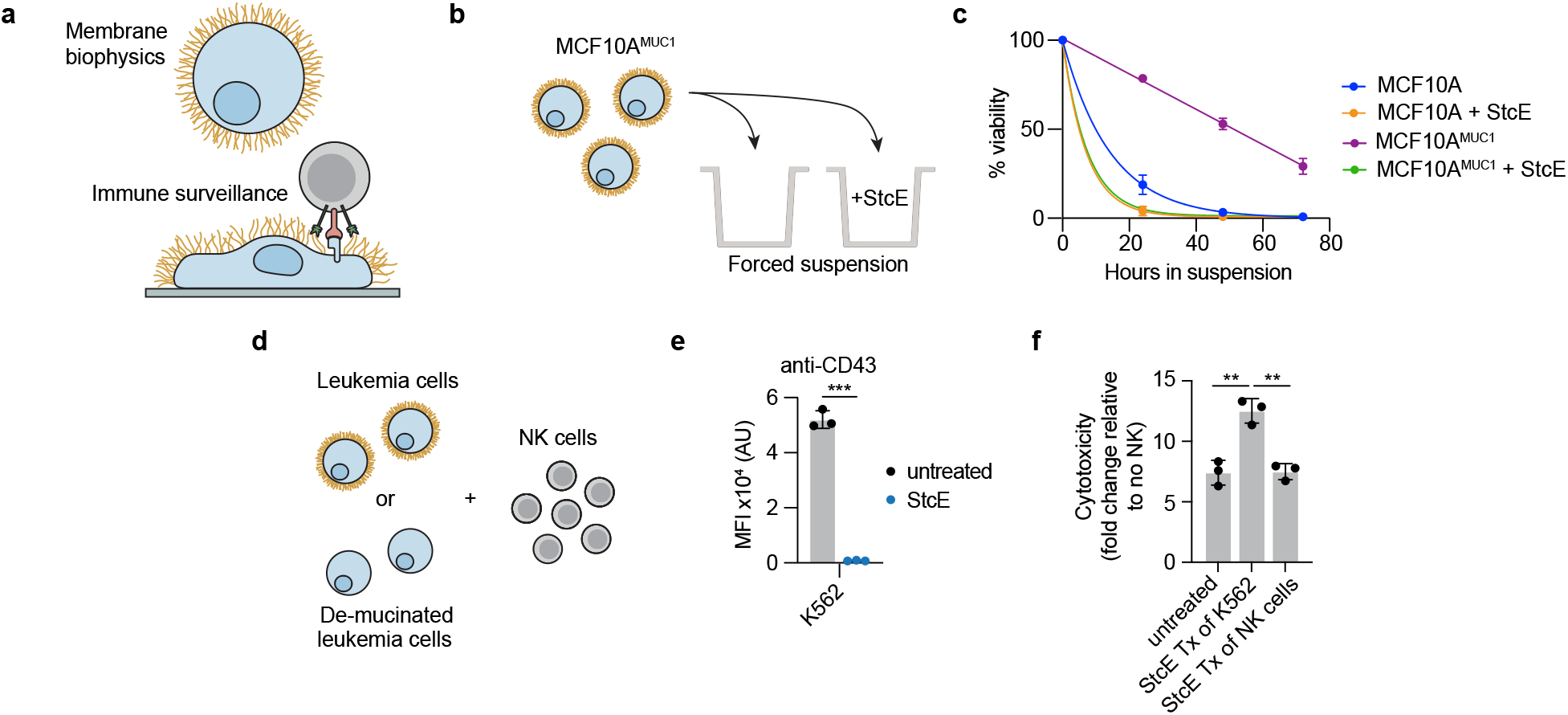
Mucinase treatment reverses mucin-driven survival pathways in cancer cell lines. **a**, Schematic depicting that mucins influence membrane biophysics and immune surveillance. **b**, Setup for suspension survival assay under anchorage-free conditions using MCF10A cells expressing doxycycline-inducible MUC1 ectodomain and treated with or without StcE mucinase. **c**, Viability of MCF10A^±MUC1^ cells ± 10 nM StcE over 72 hours as determined by flow cytometry (*n*=3 biological replicates). **d**, Setup for immune surveillance assay with leukemia cell lines ± StcE and primary human NK cells. **e**, Surface CD43 levels of K562 cells ± 50 nM StcE measured by flow cytometry (*n*=3 biological replicates). **f**, Normalized NK cell killing of K562 cells under indicated StcE treatment (Tx) conditions at a 2:1 effector:target ratio (*n*=3 biological replicates). Data are mean ± s.e.m. (**c**) or mean ± s.d. (**e-f**). *P*-values were determined using Tukey-corrected two-way ANOVA. \**p* < 0.05, \*\**p* < 0.005, \*\*\**p* < 0.0005.

Cancers, and especially carcinomas, hijack mucin signaling pathways to protect themselves from both biophysical and immunological insults. It is estimated that just one member of the mucin family, MUC1, is aberrantly expressed in greater than half of carcinomas diagnosed per year in the U.S.^6^, a frequency matched by prototypical oncogenes such as *RAS* and *MYC*. In addition, common carcinomas such as breast, ovarian, and intestinal cancers have mucinous forms, wherein tumor cells present as individual colonies suspended in a matrix of secreted mucin and polysaccharides^7^. Decades of functional, genetic, and preclinical data support depletion of cancer-associated mucins as a strategy to reverse tumor aggressiveness in a range of carcinomas^8^.

Mucins have, however, remained canonically undruggable. Therapeutic interventions face the challenge that mucin signaling occurs through the cooperative action of hundreds of arrayed epitopes and a unique, scaffolding secondary structure. There is no catalytic site to inhibit with a small molecule, nor is there a discrete functional extracellular epitope amenable to blocking with an antibody. Indeed, therapeutic depletion of cellular mucins has only been achieved in the context of Mucin 1 kidney disease, wherein a frameshifted and truncated form of MUC1 accumulates in early secretory compartments. This intracellular accumulation can be reversed with a small molecule that binds a cargo receptor, TMED9^9^.

Targeted protein degradation (TPD) has emerged as a powerful technique to address canonically undruggable targets. Classically, TPD employs bispecific molecules to traffic unwanted proteins to endogenous cellular proteolytic machinery for degradation. An advantage of this approach is that the aberrant protein is deleted, meaning the full range of its pleiotropic effects on cell signaling are reversed^10^. As conventional TPD relies on proteasomal degradation, it is limited to targets that (i) can be bound with a bridging molecule that recruits endogenous degradation machinery, and (ii) contain cytosolic domains. Recently, cytosolic delivery of an exogenous, target-selective protease achieved proteolytic manipulation of cytosolic proteins without the need to recruit proteasome-shuttling pathways^11^, and leveraging of lysosomal- shuttling receptors has enabled TPD of cell surface and secreted proteins^12^.

In order to degrade cancer-associated mucins, we developed a strategy for degradation of cell surface proteins on specific cells without the need for endogenous degradation machinery.

Specifically, a protease with dual glycan- and peptide-based selectivity for mucins is targeted to cancer cells via fusion to a nanobody. We demonstrate that these targeted proteases reduce cancer cell viability *in vitro*, and blunt primary tumor burden and metastatic outgrowth in murine breast cancer models. As nearly all extracellular proteins are glycosylated^13^, and glycosylation status is commonly altered in disease^4^, glycoform-dependent and cell type-selective TPD presents a general opportunity for increasing on-target specificity for disease-driving extracellular proteins.

## Results

### Mucinase treatment undermines mucin-driven survival pathways in cancer cells

We and others have characterized proteases from the bacterial kingdom with selectivity for mucin domains^14–18^. These “mucinases” act through recognition of joint peptide- and glycan- motifs, which have been mapped using mass spectrometry of cleavage products. As an initial candidate for therapeutic repurposing, we chose the zinc metalloprotease StcE from *Escherichia coli* serotype O157:H7. StcE recognizes the motif S/T*-X-S/T, where the first Ser/Thr must bear an O-glycan (asterisk) in order for cleavage to occur^15^. StcE is agnostic to the structure of the glycan and the identity of the X amino acid, which can also be absent. StcE is therefore a pan mucinase, able to act upon epitopes present across the natural mucins.

To begin, we tested whether treatment with StcE could reduce cell viability by undermining the biophysical function of mucins. Expression of the MUC1 ectodomain in mammary epithelial cells induces a bulky glycocalyx, which causes the cells to lift from their basement membrane and thrive in suspension in a manner characteristic of circulating metastatic tumor cells^19^. To model suspension survival *in vitro*, wild-type cells and cells overexpressing MUC1 were plated on ultralow attachment plates, treated with or without StcE, and analyzed by flow cytometry to assess viability over three days (Fig. 1b). Under these anchorage-free conditions, StcE treatment resulted in rapid cell death, consistent with previously reported alterations in membrane biophysical signaling through PI3K-Akt (Fig. 1c)^20^. Meanwhile, StcE treatment of MUC1-expressing cells in standard tissue culture plates caused suspended cells to settle, after which they continued to divide, highlighting the low toxicity of mucinase treatment at nanomolar doses (Supplemental Videos 1-2).

Next, we asked whether StcE treatment could enhance immune surveillance of cancer cells. The mucin CD43 has been recently identified as a ligand on leukemia cells for the NK cell immune checkpoint receptor Siglec-7^5^. In this model, removal of CD43 potentiates NK cell killing of leukemia cell lines. To assess whether mucinase treatment would have a similar effect, we treated three leukemia cell lines with or without endotoxin-free StcE (*Methods*), incubated them with healthy human blood donor NK cells, and quantified viability after 4 hours (Fig. 1d). StcE treatment resulted in loss of cell surface CD43 and overall Siglec-7 ligand residency, as expected (Fig. 1e and Fig. S1a-b)^5^. De-mucinated leukemia cells were susceptible to increased NK cell surveillance, consistent with a recent report showing augmentation of breast cancer line killing by immortalized NK cells upon mucinase treatment (Fig. 1f and Fig. S1c-d)^21^.

As the presence of mucins on the cell surface has been associated with decreased drug efficacy^22^, we also explored whether mucinase treatment would synergize with small molecule cytotoxic drugs. Using scalable time-lapse analysis of cell death kinetics (STACK)^23^, we screened a 261 compound library in the ovarian cancer cell line OVCAR-3 (Fig. S1e). Erastin, which induces ferroptosis through inhibition of the cystine:glutamate antiporter system x_c_^−,24^ scored among the top hits for compound cell death enhanced by StcE treatment (Fig. S1f-g and Supplemental Table 1). A dose response with erastin2, a more potent analogue, confirmed enhancement of ferroptosis with StcE co-treatment that was fully suppressed by the ferroptosis inhibitor, ferrostatin-1. In contrast, there was no enhancement of ferroptosis induced by the mechanistically distinct compound RSL3 (Fig. S1h)^25^. Taken together, these results demonstrate that removal of mucins via mucinase treatment can reverse their pleotropic tumor- progressive roles.

### Toxicity profile of StcE necessitates tumor-targeting

Bacterial enzymes are currently employed as frontline cancer therapeutics; for example, L-asparaginase from *E. coli* is used in childhood acute lymphoblastic leukemias^26^. As mucinases had not, to our knowledge, been tested as injectable therapeutics, we assayed StcE for activity and tolerability *in vivo*. The maximum tolerated dose for StcE treatment in BALB/c and C57BL/6 mice was 0.25 mg/kg. Necropsy and complete blood count (CBC) analyses performed 3 hours post injection of 15 mg/kg StcE revealed hemorrhages underneath the skull, ecchymoses throughout the gastrointestinal tract, neutrophil accumulation in the lungs, and platelet depletion (Fig. S2a-b). Western blot using a mucin-specific probe^16^ showed that StcE injected at 0.25 mg/kg remained in circulation for at least a day and digested mucins throughout the body, though not as completely as higher doses (Fig. S2c-e). As endothelial and white blood cell surface mucins are critical components of clotting and immune activation pathways^27^, these findings established that an engineered mucinase variant with selectivity for tumor-associated mucins was necessary to avoid off-target effects.

The clinical success of antibody-drug conjugates has shown that fusion of toxic therapeutic cargo to antibodies is a viable strategy for lowering off-target toxicity and increasing on-target efficacy^28^. More recently, antibody-enzyme conjugates have been designed to target the hydrolytic activity of an enzyme to specific subsets of cells^29^. An important design principle of antibody-enzyme conjugates is to ensure that the activity of the enzyme is sufficiently low such that hydrolysis only occurs when the enzyme is concentrated at its target via binding of the antibody. Specifically, in previous work with an antibody of nanomolar affinity, micromolar enzymatic activity was shown to be effective for cell surface targets^30^. As StcE is active at sub-nanomolar concentrations, our initial aim was to engineer a mucinase which retained its peptide and glycan specificity but exhibited activity within the micromolar range.

### Structure-guided engineering reduces activity, binding, and size of StcE

We turned to a previously published crystal structure^31^ to rationally design a StcE mutant with reduced activity and cell surface binding but retained specificity for mucins. To begin, we deleted two domains, the C and INS domains (Fig. 2a, *left*), as removal of the INS domain had previously been observed to reduce enzymatic activity and removal of the C domain had been shown to decrease nonspecific cell surface binding^31^, both of which were desirable for our targeted mucinase. We also mutated residues near the active site (Trp366, His367, and Tyr457), hypothesizing that these mutations would reduce activity without fully ablating catalysis or affecting mucin specificity (Fig. 2a, *right*).

**Figure 2.**
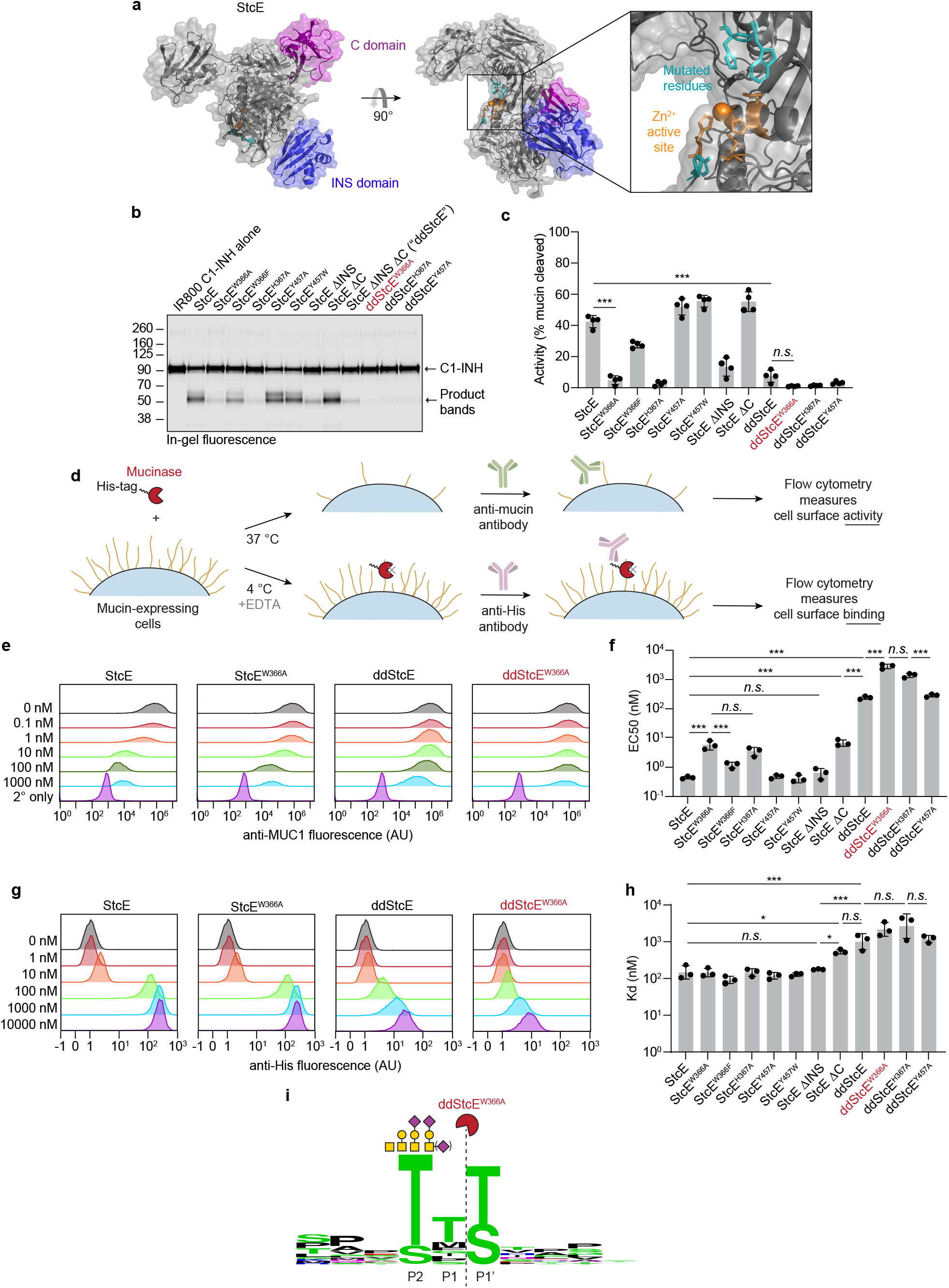
Structure-guided engineering of StcE yields mutants of reduced activity, binding, and size. **a**, Structure of StcE, as predicted by AlphaFold (*Methods*)^54^, with the C domain (purple) and INS domain (blue) highlighted. The Zn^2+^ active site is depicted in orange while mutated residues are shown in teal. **b**, Digestion of purified human mucin C1-INH with 50 nM StcE or StcE mutants, quantified by in-gel fluorescence. **c**, Quantification of (**b**) (*n*=4 independent digestions). **d**, Setup for flow cytometry assays measuring cell surface activity and binding of StcE and StcE mutants. **e**, Representative flow plots showing surface MUC1 levels of HeLa cells treated with StcE mutants at indicated concentrations. For flow plots of all other StcE mutants, see Fig. S3c. **f**, EC50 values derived from (**e**) and Fig. S3c (*n*=3 biological replicates). For dose response curves see Fig. S3d. **g**, Representative flow plots depicting cell surface binding of StcE variants on HeLa cells measured by anti-His staining. For flow plots of all other StcE mutants, see Fig. S3e. **h**, Kd values derived from (**g**) and Fig. S3e (*n*=3 biological replicates). For dose response curves see Fig. S3f. **i**, ddStcE^W366A^ cleavage motif as determined by mass spectrometry on recombinant mucins. Data are mean ± s.d. *P*-values were determined using Tukey-corrected one-way ANOVA. \**p* < 0.05, \*\**p* < 0.005, \*\*\**p* < 0.0005.

Combinations of domain deletions and single point mutations yielded a total of eleven mutants for characterization, with expression yields ranging from 20-100 mg/L (*Methods* and Fig. S3a). *In vitro* activity against recombinant mucin substrates was quantified by densitometry following SDS-PAGE (Fig. 2b-c and Fig. S3b). To determine the half maximal effective concentration (EC50) of the mutants for degradation of mucins on live cell surfaces, cells were treated with various concentrations of enzymes for 1 hour at 37 °C and subjected to flow cytometry to assess cell surface MUC1 staining (Fig. 2d-f and Fig. S3c-d). To determine effective dissociation constants (Kd) of the mutants, cells were treated with the same concentrations of enzymes for 30 minutes at 4 °C in the presence of the metal chelator and metalloprotease inhibitor EDTA to prevent enzyme activity. In the latter case, binding was quantified via interaction of an anti-His tag antibody with the His-tagged mutants (Fig. 2d,g-h and Fig. S3e-f).

Combining the two domain deletions (ΔC and ΔINS) yielded ddStcE (double deletion StcE), which exhibited reduced activity and binding, along with a molecular weight reduction to 76 kDa relative to the 98 kDa parent enzyme. Nevertheless, ddStcE’s EC50 and Kd on cells remained in the high nanomolar range. Of the single point mutations, W366A and H367A most drastically reduced activity against recombinant and cell surface mucins (Fig. 2c,f). Adding W366A and H367A mutations to the ddStcE scaffold yielded ddStcE^W366A^ and ddStcE^H367A^, which were active in the desired micromolar range, with approximate EC50 values of ∼3 and ∼1 μM, respectively, and effective Kd values of ∼2 µM each (Fig. 2f,h). ddStcE^W366A^, which we refer to as engineered StcE or “eStcE”, was selected as the scaffold for the targeted enzyme, because it exhibited the lowest activity against cell surface MUC1. We characterized the cleavage motif of eStcE using mass spectrometry as was previously done for StcE^15^, revealing that the mutations did not alter substrate recognition (Fig. 2i, Supplemental Table 2).

### A nanobody-eStcE conjugate achieves targeted mucin degradation

We envisioned that targeting eStcE to cancer cells would reverse biophysical and immunological tumor-progressive pathways while leaving bystander cells unaffected (Fig. 3a). As immune cells bind Fc-containing biologics through the Fc receptor^32^, we created a genetic fusion to a nanobody rather than an antibody. The cell surface receptor HER2 was selected as the target antigen because it is upregulated in several carcinoma subtypes, including breast and ovarian, and is bound by a well-validated nanobody, 5F7^33^. We designed two different fusion orientations which we tested for expression yield, stability, mucinase activity, and cell surface HER2 binding (Fig. 3b). Both orientations of the chimera expressed in endotoxin-deficient ClearColi at 30-60 mg/L and were similarly active against recombinant MUC16 as eStcE alone (Fig. S4a). The conjugate with the mucinase C-terminal to the nanobody, which we refer to as “αHER2-eStcE”, was stable and retained activity after months at 4 °C, while the other orientation “eStcE-αHER2” exhibited reduced activity following equivalent storage conditions (Fig. S4b). Effective dissociation constants (Kd) for nanobody-mucinase conjugates were determined via flow cytometry of HER2+ cells as described above, giving values of 11, 4, and 58 nM for αHER2, αHER2-eStcE, and eStcE-αHER2, respectively (Fig. S4c-e). Therefore, αHER2-eStcE was selected for further *in cellulo* and *in vivo* analyses due to its increased stability and binding to HER2+ cells.

**Figure 3.**
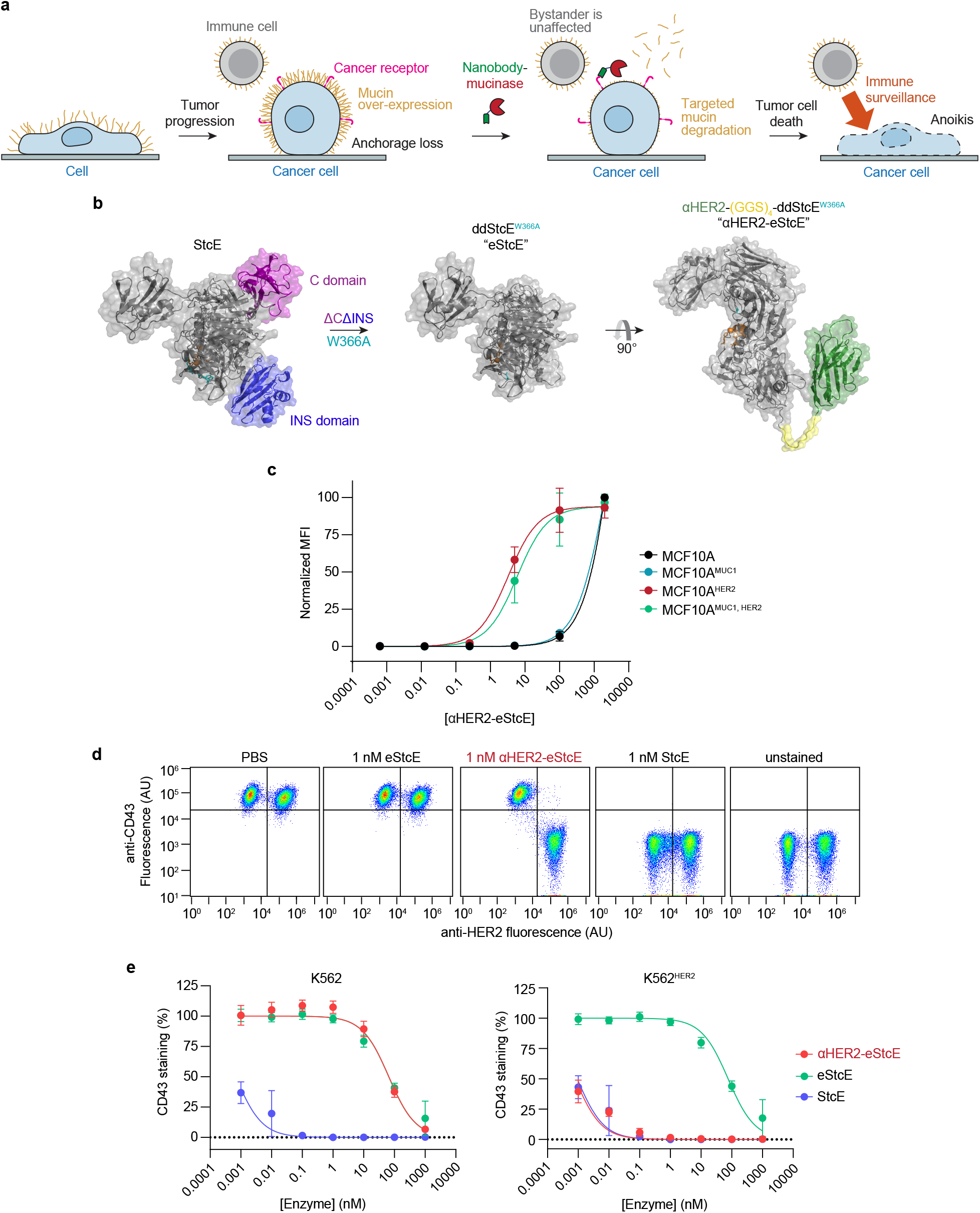
An optimized nanobody-mucinase conjugate selectively cleaves mucins from HER2+ cells. **a**, Schematic depicting reversal of mucin-driven tumor progressive pathways via treatment with a targeted nanobody-mucinase conjugate. **b**, Structure of nanobody-mucinase conjugate, as predicted by AlphaFold (*Methods*)^54^, with engineering strategy shown. The HER2-targeting nanobody is depicted in green, active site is shown in orange, mutated residue (W366A) in teal, and flexible linker in yellow. **c**, Binding curves of αHER2-eStcE on MCF10A^±MUC1, ±HER2^ cells measured by anti-His staining as in Fig. 2d (*n*=3 biological replicates). For flow plots and Kds see Fig. S4f-g. **d**, Representative flow plots depicting surface CD43 levels of mixed K562^±HER2^ cells treated with 1 nM mucinases or conjugate overnight. For representative flow plots with shorter incubation times see Fig. S6e. **e**, CD43 cleavage curves derived from (**d**) (*n*=3 biological replicates). Data are mean ± s.d.

We used cells transduced with a doxycycline-inducible MUC1 ectodomain construct to test the contributions of the nanobody and the enzyme components of αHER2-eStcE to mucin binding. αHER2-eStcE bound to HER2+ cell surfaces with a Kd value approximately three orders of magnitude higher relative to its binding to HER2- cells, indicating that αHER2-eStcE bound to cells via HER2 affinity and not mucin affinity (Fig. 3c and Fig. S4f-g). Cleavage assays with a panel of recombinant non-mucin and mucin proteins confirmed that the fusions maintained mucin selectivity (Fig. S5a). In order to test the specificity of αHER2-eStcE for mucins versus non-mucin proteins on cell surfaces, we employed terminal amine isotopic labeling of substrates mass spectrometry (TAILS MS), which is a method optimized for detection of peptides generated from protease digestion of live cells^34^. A HER2+ suspension cell line was treated with vehicle control, StcE, eStcE, or αHER2-eStcE, and supernatants were collected and subjected to TAILS MS (Fig. S5b). Importantly, analysis of peptides generated relative to vehicle control confirmed selectivity for mucin-domain glycoproteins, similar cleavage profiles between StcE and αHER2-eStcE, and reduced activity for untargeted eStcE (Fig. S5c-f).

Next, we tested the conjugate’s on-target activity through mixed cell assays with various human cancer cell lines engineered to express HER2 (Fig. S6a-d). Mixed HER2+ and HER2- cells were treated with StcE, eStcE, or αHER2-eStcE overnight, and depletion of cell surface mucins was analyzed via live cell flow cytometry to quantify anti-CD43 staining. StcE treatment at 1 nM resulted in complete removal of cell surface mucins on both HER2+ and HER2- cells, while 1 nM of eStcE resulted in no discernable removal of mucins in either population. In contrast, 1 nM of αHER2-eStcE resulted in complete loss of cell surface mucins on HER2+ cells and no discernable loss of mucins on HER2- cells (Fig. 3d, quantified in Fig. 3e, time course in Fig. S6e). The same trend was observed at higher doses in another cell line interrogated for cell surface residency of a different mucin protein, MUC1 (Fig. S6f-g).

### αHER2-eStcE selectively kills HER2+ cells in mixed cell assays and is nontoxic in mice

Using a series of mixed cell assays, we tested whether αHER2-eStcE could selectively reverse mucin-dependent tumor-progressive pathways. We repeated biophysical and immunological assays from Figure 1 using HER2+ and HER2- cell populations, which were mixed prior to enzymatic treatment (Fig. 4a,c). Under anchorage-free conditions, αHER2-eStcE reversed suspension survival in only the HER2+ population whereas StcE treatment resulted in cell death in both populations (Fig. 4b). Likewise, treatment with αHER2-eStcE selectively enhanced NK cell-mediated killing of the HER2+ population (Fig. 4d). Recently, it was found that removal of cell surface mucins via mucinase treatment promotes engagement of trans-acting phagocytic receptors in macrophages^35^. Thus, we tested αHER2-eStcE in a mixed cell assay with primary human macrophages and immortalized human breast cancer cells, where we observed enhancement of phagocytosis of HER2+ cells relative to HER2- cells (Fig. S6h-j).

**Figure 4.**
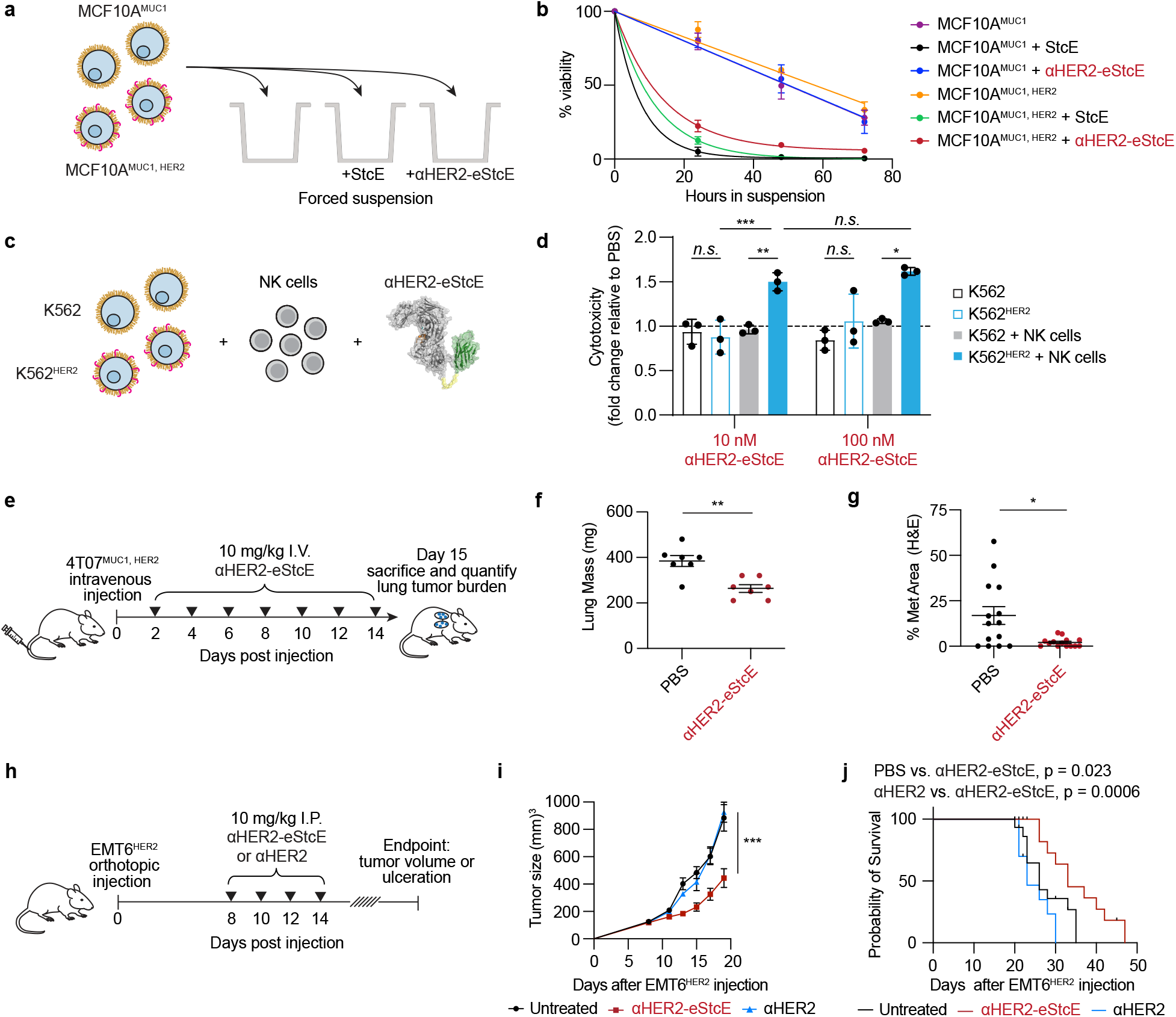
αHER2-eStcE is effective in mixed cell assays and breast cancer mouse models. **a**, Setup for mixed cell suspension survival assay under anchorage-free conditions as in Fig. 1b. **b**, Viability of mixed MCF10A^MUC1, ±HER2^ cells ± 1 nM StcE or αHER2-eStcE over 72 hours as determined by flow cytometry (*n*=3 biological replicates). **c**, Setup for mixed cell NK cell killing as in Fig. 1d. **d**, Normalized NK cell killing of mixed K562^±HER2^ cells treated with conjugate at a 2:1 effector:target ratio (*n*=3 biological replicates). **e**, Treatment regimen for BALB/c mice injected intravenously (I.V.) via tail vein with 4T07^MUC1, HER2^ cells. αHER2-eStcE was injected I.V. every other day at 10 mg/kg starting on day 2 (*n*=7 animals per group). **f**, Plot depicting lung masses of animals described in (**e**). **g**, Percent area of lung metastases quantified by H&E tissue staining of animals described in (**e**). Points represent individual scans of lung slices (*n*=2 per animal). For images, see Fig. S9d- e. **h**, Treatment regimen for BALB/c mice injected with EMT6^HER2^ orthotopically into the mammary fat pad. αHER2-eStcE or αHER2 were injected four times intraperitoneally (I.P.) every other day starting on day 8 (*n*=9-12 animals per group). The dose was 10 mg/kg for αHER2-eStcE and an equimolar quantity (2.8 nmol) of αHER2. **i**, Average growth curves of EMT6^HER2^ tumors for animals described in (**h**). **j**, Survival curves for animals described in (**h-i**). Data are mean ± s.e.m. (**b, f-g, i**) or mean ± s.d. (**d**). *P*-values were determined using Tukey- corrected two-way ANOVA and Mantel-Cox test. *p < 0.05, **p < 0.005, ***p < 0.0005.

Intravenous administration of fluorophore-labeled αHER2-eStcE at doses ranging from 0.25-10 mg/kg into BALB/c mice revealed that the conjugate remained in blood and tissues for approximately 24 hours, with no discernable toxicity (Fig. S7a-b). Blinded necropsy and complete blood count analyses confirmed no abnormalities at the highest tested dose of 10 mg/kg (Fig. S7c-d). To assess mucin depletion in tissues, we injected 5 mg/kg StcE or αHER2- eStcE intravenously into BALB/c mice, collected plasma, liver, spleen, and lung 4 hours post injection, and immunoblotted for mucins. In all cases, αHER2-eStcE injection resulted in significantly reduced mucin depletion when compared to the wild-type parent enzyme (Fig. S7e).

### αHER2-eStcE blunts primary tumor burden and metastatic outgrowth in models of murine breast cancer

To assess on-target efficacy of αHER2-eStcE we turned to previously validated murine models of breast cancer progression. The murine cell line 4T07 is a BALB/c syngeneic mammary carcinoma that efficiently metastasizes to sites such as the lung, but is unable to efficiently proliferate at metastatic sites^36^. Woods *et al*. showed that elaboration of 4T07 cell surfaces via ectopic expression of MUC1 ectodomain or with lipid-anchored mucin mimetic glycopolymers enhances proliferation in the metastatic niche through PI3K-Akt mechanosignaling pathways related to cell cycle progression^20^. This model involved tail vein injection of luciferase-expressing 4T07 cells into BALB/c mice, whereupon cells were lodged in the small capillaries of the lung. At day 15 post injection, animals were sacrificed and tumor burden in the lung was quantified by lung mass and immunohistochemistry.

We performed a therapeutic model with 4T07 cells stably expressing MUC1 ectodomain and HER2, to assess whether αHER2-eStcE would influence metastatic outgrowth to the lung. Treatment was administered intravenously every other day with 10 mg/kg αHER2-eStcE or vehicle control (Fig. 4e). The dosing strategy was chosen based on (i) the observed 24-hour *in vivo* circulation time (Fig. S7a), and (ii) the approximately 24-hour turnover observed *in cellulo* for enzymatically degraded mucins (Fig. S8), consistent with reported mucin half-lives^37^. Bioluminescent imaging (BLI) directly following injection confirmed 4T07 cells seeded lungs of both control and treatment group animals (Fig. S9a-b). BLI at 13 days post injection revealed reduced tumor burden in αHER2-eStcE treated animals (n = 7, p = 0.097). Total mouse mass did not differ significantly between groups (Fig. S9c). Necropsy of animals on day 15 post injection showed a decreased wet lung mass (n = 7, p = 0.0041) and decreased percent metastatic area by H&E tissue staining (n = 7 mice, two lung sections per mouse, p = 0.0061) (Fig. 4f-g and Fig. S9d-e). Immunohistochemistry (IHC) analysis of lungs revealed reduction in pFAK-Y397 and cyclin D1 staining in αHER2-eStcE treated versus control animals (Fig. S10- 11). These data indicate that treatment with αHER2-eStcE may blunt metastatic outgrowth in the 4T07 model through reversal of mucin ectodomain-driven enhancement of cell cycle progression via the PI3K-Akt axis^20^.

The murine cell line EMT6 is a BALB/c syngeneic mammary carcinoma that is used as a model for immune surveillance^38^. Gray *et al*. showed that desialylation of orthotopic EMT6 tumors with injected sialidase constructs prolonged the survival of mice through inhibition of the Siglec-sialic acid immune checkpoint axis^30^. This model involved injection of EMT6 cells stably expressing HER2 (EMT6^HER2^) into the mammary fat pads of mice followed by I.P. treatment with enzymes or controls. Tumor size was measured with calipers until tumor burden required euthanasia (typically 20-30 days post injection).

To assess whether mucinase-driven depletion of Siglec ligands from tumor cell surfaces would have a similar beneficial effect, we performed a therapeutic model with EMT6^HER2^ cells and treated animals with four doses of 10 mg/kg αHER2-eStcE, αHER2, or vehicle control (Fig. 4h). Treatment with αHER2-eStcE resulted in reduced tumor size at day 19 (n = 6-9, p < 0.0001) (Fig. 4i). After 35 days, all mice in the vehicle-treated group had reached a tumor burden requiring euthanasia, while treatment with αHER2-eStcE extended mouse survival to 47 days (p = 0.023 versus control and p = 0.0006 versus αHER2 alone) (Fig. 4j). Treatment with αHER2 alone did not result in attenuation of tumor growth or prolonged survival. Mice treated with αHER2-eStcE did not exhibit weight loss over the course of the experiment, suggesting that treatment was well tolerated. (Fig. S12a).

For analysis of mucin degradation and immune infiltration within EMT6^HER2^ tumors, a separate set of animals were treated as above with vehicle, αHER2, or αHER2-eStcE, and sacrificed at day 10 post-implantation (Fig. S12b). Importantly, αHER2-eStcE degraded mucins on the EMT6^HER2^ cancer cells (CD45^−^/HER2^+^ cells) but did not affect mucin levels on immune cells (CD45^+^/HER2^−^ cells), indicating αHER2-eStcE promotes selective mucin depletion *in vivo* (Fig. S12c-e).

We next profiled the immune composition within EMT6^HER2^ tumors and found that the dominant immune cell type within these tumors were Ly6G+ cells, which correspond to Ly6G-expressing granulocytes and/or neutrophils that are often found in breast tumor immune infiltrates (Fig. S13a-d)^39^. Tumor-infiltrating Ly6G+ cells from mice treated with αHER2-eStcE showed reduced levels of the inhibitory immune checkpoint PD-1 relative to vehicle and αHER2 treatment groups (Fig. S13e-f). In addition, αHER2-eStcE therapy promoted infiltration of conventional dendritic cells (cDCs) into the tumors (Fig. S13g). We found that cDCs in αHER2-eStcE treated tumors exhibited an augmented phenotype, as indicated by reduced levels of the inhibitory ligand PD- L1 (Fig. S13i-j). Strikingly, cDCs of conjugate treated animals also exhibited significantly increased levels of granzyme B (GzmB), a cytotoxic protease that is released by immune cells to trigger apoptosis of target cells (Fig. S13h). While GzmB is typically associated with cytotoxic cells, such as CD8^+^ T cells and NK cells, it can also be produced by other cell types upon activation^40^. These data, which support modulation of the tumor immune microenvironment following αHER2-eStcE treatment, are consistent with the importance of mucin and sialic acid signaling in the tumor microenvironment^41–43^.

In summary, targeted degradation of cancer-associated cell surface mucins limited primary tumor growth and metastasis *in vivo* in murine models of breast cancer progression, with indications that both biophysical and immunological signaling were influenced.

## Discussion

Over 80% of cell surface and secreted proteins are predicted to be glycosylated^13^, and specific alterations in protein glycosylation have been observed in diverse pathologies including inflammation and cancer^4^. An emerging theme from research in this field is that the biological functions of these altered glycoproteins are determined equally by their glycan and protein components. Accordingly, specific disease-relevant protein glycoforms are an important class of therapeutic targets^4^. This realization motivated us to consider new therapeutic modalities that target composite epitopes comprising both protein and glycan. Mucin glycoproteins are attractive targets for this approach since their structures and biological activities are equally dependent on their protein and glycan constituents. We and others have characterized bacterial proteases with peptide- and glycan-dependent cleavage motifs that render them highly selective for densely O-glycosylated mucin domains^14–18^. Therefore, we chose mucinases, specifically the pan mucinase StcE, as an initial candidate for evaluation as a peptide- and glycan-selective mucin degrader.

Antibody-enzyme conjugates have been previously used to spatially restrict enzymatic activity in therapeutic contexts. In one technique, termed antibody directed enzyme prodrug therapy (ADEPT), the enzyme activates a prodrug, resulting in selective release of active compound at a tumor or pathogen site^44^. In another strategy, the enzyme is hydrolytic, resulting in degradation of the antibody’s target or cell death in the vicinity of the affinity reagent’s binding site^45^. Building upon this prior work, we recently reported antibody-sialidase conjugates for selective desialylation of tumor cell surfaces, which have entered Phase I/II clinical trials (trial identifier NCT05259696)^29,30^.

To design antibody-mucinase conjugates with high on-target selectivity we drew from insights gained through efforts to engineer so-called molecular glues, which are small molecules that bridge a target protein with an endogenous enzyme^46,47^. Molecular glues, such as proteolysis targeting chimeras (PROTACs), operate through the principle of effective molarity, wherein two reactants that are in close proximity interact pseudo-intramolecularly, dramatically increasing the reaction rate^48^. Selectivity against a target pool is achieved when two criteria are met: (i) binding of the enzyme conjugate to its substrates (measured for example by Kd) is driven by the affinity of the targeting moiety and not the enzyme’s intrinsic affinity, and (ii) the enzyme’s activity (measured for example by EC50) is such that intermolecular reactions occur slowly relative to pseudo-intramolecular reactions. Based upon prior work with antibody-sialidase conjugates, we reasoned both criteria could be met with a high affinity antibody and a low efficiency enzyme^30^.

A series of domain deletions and point mutations yielded an engineered StcE variant which we term eStcE, exhibiting micromolar Kd and EC50 values on cell surfaces. We fused eStcE to an anti-HER2 nanobody (αHER2), which binds the human HER2 receptor with nanomolar affinity^33^. We expected the lack of an immune cell binding Fc domain on αHER2 to be an advantage, given widespread expression of functionally important mucins such as CD45 and MUC1 on immune cell surfaces^27^. The conjugate, termed αHER2-eStcE, exhibited high on-target activity as quantified by mixed cell assays in a variety of cell types expressing different mucin proteins. In cell lines, functional mixed cell assays replicated the biophysical and immunological effects of treatment with the wild-type StcE enzyme and showed high on-target selectivity. In a mouse model of mechanical signaling during lung metastatic outgrowth, αHER2-eStcE treatment decreased lung burden in mucin-overexpressing tumors^20^. In an orthotopic mouse model of breast cancer, previously shown to be driven by sialic acid-mediated immunological silencing^30^, αHER2-eStcE treatment decreased tumor burden and increased survival.

Future work will focus on determining the precise therapeutic contexts in which targeted mucin degradation would be most effective. Based on our data, tumors which are immune-infiltrated, susceptible to mechanical stress, and sensitive to ferroptosis may be a starting point. Cancers which present as so-called “mucinous” subtypes, wherein individual cancer cells are suspended in a secreted matrix of mucin and polysaccharides, may be a second set of indications^7^. In addition, the range of human diseases which are characterized by aberrant mucin phenotypes, such as respiratory viral infections^49,50^, cystic fibrosis^51^, bacterial endocarditis^52^, and gut dysbiosis^53^, are a third class of disease indications. Further experiments in preclinical models will be necessary to assess whether nanobody-mucinase conjugates have the potential to join other bacterially-derived enzymes, such as asparaginase^26^, as frontline therapeutics. Finally, identification of human proteases capable of degrading mucin domains would be of benefit to clinical translation of this class of biologics.

## Acknowledgements

This work was supported, in part, by National Cancer Institute Grant R01CA227942 (to C.R.B.) and Swiss National Science Foundation (SNSF Nr. 310030_184720/1 (to H.L.). K.P. was supported by a U.S. National Science Foundation Graduate Research Fellowship, a Stanford Graduate Fellowship, and the Stanford ChEM-H Chemistry/Biology Interface Predoctoral Training Program. D.J.S. was supported by a U.S. National Science Foundation Graduate Research Fellowship and Stanford Graduate Fellowship. G.S.T. was supported by a Hertz Foundation Fellowship, a U.S. National Science Foundation Graduate Research Fellowship, and the Stanford ChEM-H Chemistry/Biology Interface Predoctoral Training Program. K.J.M. was supported by an American Cancer Society–2017 Seattle Gala Paddle Raise Postdoctoral Fellowship (PF-18-118-01-CDD) and the Program for Breakthrough Biomedical Research, which is partially funded by the Sandler Foundation. S.W. was supported by a Banting Postdoctoral Fellowship from the Canadian Institutes of Health Research (CIHR). N.M.R. acknowledges support from NIH/NCI K00 Predoctoral to Postdoctoral Transition Award K00CA212454. A.K. and C.L.M. were supported by U.S. National Science Foundation Graduate Research Fellowships.

## Author contributions

K.P., D.J.S., G.S.T., N.R.M., J.J.N., K.J.M., S.P.W., N.M.R., G.C.F., A.K., and B.M.G. designed and carried out experiments. S.A.M. performed mass spectrometry and data analysis for conjugate cleavage motif. K.M.C. and J.G.V.-M. performed complete necropsies and histopathology for toxicity experiments. C.L.M. contributed to data analysis. H.L., V.M.W., and C.R.B. provided conceptualization and supervision. K.P., D.J.S., G.S.T., and C.R.B. wrote the manuscript with input from all authors.

