## Supplemental Information for "Design of a mucin-selective protease for targeted degradation of cancer-associated mucins"

**This PDF file includes:**

Materials and Methods
Figs. S1 to S14
Tables S3 to S5

**Other Supplementary Materials for this manuscript include the following:**

Movies S1 to S2
Tables S1 to S2

### Materials and Methods

**Cell culture.** Cells were maintained at 37 ºC, 5% $CO_2$. MCF10A$^{\pm MUC1, \pm HER2}$ cells were cultured in phenol red free 1:1 DMEM:F12 supplemented with 5% New Zealand horse serum (Thermo Fisher Scientific), 20 ng/mL epidermal growth factor (Peprotech), 0.5 µg/mL hydrocortisone (Millipore Sigma), 100 ng/mL cholera toxin (Millipore Sigma), 10 µg/mL insulin (Millipore Sigma), and 1% penicillin/streptomycin (P/S). K562$^{\pm HER2}$, CCRF-CEM, and 4TO7$^{MUC1, HER2}$ cells were cultured in RPMI supplemented with 10% heat inactivated fetal bovine serum (FBS) (Thermo Fisher Scientific) and 1% P/S. HeLa, CCRF-HSB-2, EMT6$^{HER2}$, and HEK-293T cells were grown in DMEM supplemented with 10% heat inactivated FBS and 1% P/S. MCF7 cells were grown in DMEM supplemented with 10% heat inactivated FBS and 1% P/S and 10 µg/mL human insulin (Thermo Fisher Scientific). OVCAR-3$^{N}$ cells were cultured in RPMI supplemented with 10% heat inactivated FBS, 0.01 mg/mL bovine insulin (Sigma-Aldrich), and 1% P/S. Cells were counted using Countess II FL Automated Cell Counter (Thermo Fisher Scientific) following manufacturer's recommendations.

**Supplementary Table 3. Antibody clones and working concentrations.**

| Antibody | Vendor, clone | Dilution/concentration |
|---|---|---|
| AF488/647 anti-human CD340 (erbB2/HER-2) antibody | BioLegend, 24D2 | 1:200 |
| AF647 CD43/sialophorin antibody | Novus Biologicals, MEM-59 | 1:1000 |
| Anti-MUC1/episialin antibody | EMD Millipore, 214D4 | 1:200 (flow cytometry), 1:1000 (Western blot) |
| AF647 Affinipure Goat Anti-Mouse IgG | Jackson ImmunoResearch | 1:375 |
| FITC anti-His antibody | Miltenyi Biotec, GG11-8F3.5.1 | 1:50 |
| MUC1 mouse mAb | Cell Signaling Technology, VU4H5 | 1:200 |
| MUC16 mouse mAb | Abcam, X75 | 1:1000 |
| *InVivo*MAb anti-mouse/human/rat CD47 | BioXCell, MIAP410 | 20 µg/mL |
| BV421 anti-mouse CD45 antibody | BioLegend, 30-F11 | 1:80 |
| Purified Rat Anti-Mouse CD16/CD32 (Mouse BD Fc Block) | Clone 2.4G2 | 1:50 |
| Anti-phospho-FAK tyrosine 397 | Invitrogen, Cat. #: 700255 | 1:100 |
| Anti-Cyclin D1 | Cell Signaling Technology, Cat. #: 2922 | 1:200 |

**Supplementary Table 4. Flow panel antibodies related to Fig. S13-14.**

| Antibody | Color | Vendor, Cat. No. | Cat. Number | Dilution |
|---|---|---|---|---|
| CD45 | BUV395 | BD Biosciences | 564279 | 1:100 |
| Live-dead | Zombie UV | BioLegend | 423107 | 1:100 |
| CD4 | BUV496 | BD Biosciences | 93937 | 1:100 |
| Ly-6G | BUV563 | BD Biosciences | 612921 | 1:200 |
| NKp46 | BUV661 | BD Biosciences | 741678 | 1:70 |

| | | | | |
|---|---|---|---|---|
| CD3 | BUV805 | BD Biosciences | 749276 | 1:70 |
| PD-L1 | BV421 | BioLegend | 124315 | 1:150 |
| LFA-1 | SB436 | eBioscience | 62011180 | 1:100 |
| CD8 | eFluor 450 | eBioscience | 480081 | 1:100 |
| MHCII | BV510 | BioLegend | 107635 | 1:300 |
| CD80 | BV605 | BioLegend | 104729 | 1:70 |
| CD103 | BV650 | BD Biosciences | 748256 | 1:70 |
| CD206 | BV711 | BioLegend | 141727 | 1:100 |
| PD-1 | BV785 | BioLegend | 135225 | 1:100 |
| CD19 | BB515 | BD Biosciences | 564509 | 1:100 |
| CD11c | FITC | BioLegend | 117306 | 1:100 |
| Ly-6C | PerCP | BioLegend | 128028 | 1:200 |
| Tim-3 | BB700 | BD Biosciences | 747619 | 1:100 |
| Siglec E | PE | BioLegend | 677104 | 1:50 |
| CD25 | PE-Cy5.5 | eBioscience | 35025182 | 1:100 |
| F4/80 | AF647 | BioLegend | 123122 | 1:100 |
| CD11b | APC-Cy7 | BioLegend | 101226 | 1:100 |
| Ki67 | AF532 | eBioscience | 58569882 | 1:200 |
| TCF-7 | AF700 | R&D Systems | FAB82224N | 1:100 |
| GzmB | PE-eFluor610 | eBioscience | 61889882 | 1:100 |
| FoxP3 | APC | eBioscience | 17577382 | 1:100 |

**MCF10A$^{MUC1}$ suspension survival assay.** MCF10A cells expressing a cytoplasmic truncation of MUC1 (MUC1ΔCT, also referred to as *MUC1 ectodomain*) were used to limit any possible cytoplasmic signaling[20]. MUC1ΔCT was induced with 200 ng/mL doxycycline for 24 h. Uninduced and induced cells were seeded at $3x10^5$ cells/well in a 24-well ultra-low attachment plate (Corning) in 0.75 mL of complete media. 200 ng/mL doxycycline and 10 nM StcE were added as appropriate. The plate was incubated at 37 ºC, 5% $CO_2$, 125 rpm. At t=0, 24, 48, and 72 h, cells were spun down at 350 *x g* for 5 min, resuspended in 200 µL of phosphate-buffered saline (PBS) with 0.1% benzonase (Sigma-Aldrich), and incubated at room temperature for 15 min. Cells were then resuspended in 200 µL of enzyme-free cell dissociation buffer (Thermo Fisher Scientific) and stained with 100 nM Calcein AM (Thermo Fisher Scientific) and 5 nM Sytox Red Dead Cell Stain (Thermo Fisher Scientific) for 20 min at 4 ºC, prior to analysis using a BD Accuri C6 plus. Videos of MCF10A$^{MUC1}$ cells freshly seeded on standard tissue culture plates (Corning) and treated with and without 1 nM StcE were generated with images taken at 30 min intervals for 18 h using an IncuCyte. The IncuCyte was set to 37 °C and 5% $CO_2$, and cells were incubated in complete media with 200 ng/mL doxycycline.

For mixed cell assays, MUC1ΔCT was induced with 200 ng/mL doxycycline for 24 h. A 1:1 mixture of $2.5x10^5$ MCF10A$^{MUC1}$ and MCF10A$^{MUC1, HER2}$ cells were seeded per well in a 24-well ultra-low attachment plate in 0.80 mL of complete media. 200 ng/mL doxycycline, 1 nM StcE, and 1 nM αHER2-eStcE were added as appropriate. The plate was incubated at 37 ºC, 5% $CO_2$, 125 rpm. At t=0, 24, 48, and 72 h, cells were spun down at 350 *x g* for 5 min, resuspended in

100 µL of PBS with 0.1% benzonase, and incubated at room temperature for 15 min. Cells were then resuspended in 200 µL of enzyme-free cell dissociation buffer and stained with Alexa Fluor 647 anti-human CD340 (erbB2/HER-2) antibody (24D2 clone) (BioLegend) and 500 nM Sytox Green Nucleic Acid Stain (Thermo Fisher Scientific) for 20 min at 4 ºC, according to manufacturer recommendations, prior to analysis using a BD Accuri C6 plus. All flow cytometry data were analyzed using FlowJo v. 10.0 (TreeStar).

**CD43 and Siglec-7-Fc flow cytometry.** $1x10^6$ K562, CCRF-CEM, or CCRF-HSB-2 cells growing in log phase were harvested, resuspended in 1 mL of serum-free RPMI, and treated with either vehicle or 50 nM StcE for 1 h. Cells were subsequently spun down at 600 *x g* and washed twice in PBS. Cells were then resuspended in FACS buffer (0.5% BSA in PBS) at $1x10^6$ cells/mL and aliquoted into a V-bottom 96-well plate (Corning) at $1x10^5$ cells/well. For staining, a precomplex solution of 1 µg/mL Siglec-7-Fc (R&D Systems) and 1 µg/mL Alexa Fluor 488-antihFc was made up in FACS buffer and incubated on ice for 1 h. Alexa Fluor 647 CD43/sialophorin antibody (MEM-59 clone) (Novus Biologicals) was subsequently added to the precomplex solution prior to staining. Cells were stained in 100 µL of staining solution for 30 min, washed twice with FACS buffer, and analyzed by flow cytometry using a BD Accuri C6 plus.

**Human donor-derived macrophage isolation.** Peripheral blood mononuclear cells (PBMCs) were isolated from LRS chambers (Stanford Blood Center) using a Ficoll-Paque density gradient (Cytiva). Isolated PBMCs were extracted from the PBS/Ficoll interface and washed three times with PBS. PBMCs were resuspended in RPMI containing 10% heat inactivated FBS and plated at $1x10^7$ cells/well into a 24-well #1.5 glass plate (Cellvis) that was pre-coated with poly-L-lysine solution (Millipore Sigma). PBMCs were incubated for 1 h at 37 °C to allow monocytes to adhere to the glass. Cells were then rinsed three times with PBS to remove contaminating lymphocytes. Media was replaced with IMDM containing 10% human AB serum (Gemini). Monocytes were differentiated for 7-9 days.

**NK cell isolation.** PBMC aliquots were quickly thawed and diluted in 10 mL of RPMI containing DNAse to break up cell aggregates. Cells were incubated at 5% $CO_2$, 37 °C for 30 min and subsequently counted in duplicate. Cells were then spun down at 600 *x g* and resuspended in RPMI to a final cell concentration of $50x10^6$ cells/mL. Isolation of NK cells was performed according to manufacturer's instructions using an NK cell magnetic isolation kit (STEMCELLTechnologies). NK cells were cultured for at least 24 h before conducting experiments. For killing experiments, NK cells were cultured for 24 h in complete media containing 0.2-0.5 µg/mL IL-2 (BioLegend).

**NK cell killing assays.** Target cells were harvested by centrifugation and resuspended in serum-free RPMI containing 5 µM Cell Tracker Far Red (Thermo Fisher Scientific) at $5x10^5$ cells/mL. Cells were then incubated for 30 min at 37 ºC; where indicated, cells were treated with 10-20 nM StcE. Following staining and StcE treatment, cells were spun down, washed twice with PBS containing 1 mM EDTA, and resuspended in complete media. Cells were diluted to $1x10^5$ cells/mL in complete media containing 100 nM Sytox Green, and 100 µL of cell suspension was aliquoted into a flat bottom 96-well plate. Separately, NK cells were diluted to various cell concentrations to generate the indicated effector:target ratios in complete media containing 100 nM Sytox Green. Where indicated, these cell suspensions were treated with 20 nM StcE for 30 min, washed twice with PBS containing 1 mM ETDA, and resuspended in complete media containing 100 nM Sytox Green. 100 µL of these cell suspensions was then mixed with the target cell suspensions to generate a total volume of 200 µL. Cells were incubated at 37 ºC for 4 h and analyzed by flow cytometry.

For mixed cell assays, K562$^{HER2}$ cells were harvested by centrifugation and incubated for 30 min at 37 °C in serum-free RPMI containing 0.33 μM CellTrace Far Red (Thermo Fisher Scientific) at $5x10^5$ cells/mL. K562, isolated NK, and labeled K562$^{HER2}$ cells were harvested by centrifugation and resuspended in complete media. $1x10^4$ K562 cells, $1x10^4$ K562$^{HER2}$ cells, and $2x10^4$ NK cells in 200 μL of complete media containing 50 nM Sytox Green were added to a flat bottom 96-well plate. PBS, αHER2-eStcE, or StcE in PBS were added to a final volume of 222.2 μL and incubated for 4 h at 37 °C prior to analysis by flow cytometry using a BD Accuri C6 plus.

**Bioactive compound library screen**. A library of 261 bioactive compounds (Selleck Chemicals) was stored at −80 °C. The library was re-formatted from 96-well to 384-well format using a Versette automated liquid handler configured with a 96-channel pipetting head and diluted to 2 mM in DMSO. The day before the screen, $5x10^3$ OVCAR-3$^N$ cells/well were seeded into two 384-well plates in 45 μL of media. The next day, the media was removed and replaced with media containing 20 nM Sytox Green and compounds from a freshly thawed library master stock plate (1 compound/well) were added to a final concentration of 500 nM. One plate was co-treated with vehicle (PBS) and the other with 50 nM StcE. Plates were imaged immediately and every 2 h thereafter for a total of 72 h using the Essen IncuCyte Zoom. Counts of Sytox Green and mKate2 objects per $mm^2$ were obtained and the lethal fraction calculated as previously described[23]. The area-under-the-curve (AUC) of lethal fraction scores across the full 72 h were calculated using the trapezoid rule in Excel (Microsoft Corp.). The Bliss Independence Model was used to compute expectant cell death of compound + StcE treatment using normalized AUC (nAUC) values, and deviation from this expectation was used to infer modulation of cell death, as described previously[55].

**Measuring cell death using STACK.** Follow-up cell death experiments of OVCAR-3$^N$ cells were performed using scalable time-lapse analysis of cell death kinetics (STACK)[23]. Cell lines stably expressing nuclear-localized mKate2 (Nuc::mKate2) were incubated in media containing 20 nM Sytox Green. Counts of live (mKate2$^+$) and dead (SG$^+$) objects were obtained from images collected every 2 or 4 h. The following image extraction parameter values were used to count OVCAR-3$^N$ mKate2$^+$ objects: Parameter adaptive, threshold adjustment 1; Edge split on; Edge sensitivity 50; Filter area min 20 μm$^2$, maximum 8100 μm$^2$; Eccentricity max 1.0; and SG$^+$ objects: Parameter adaptive, threshold adjustment 10; Edge split on; Edge sensitivity −5; Filter area min 20 μm$^2$, maximum 750 μm$^2$; Eccentricity max 0.9. Counts were exported to Excel and lethal fraction (LF) scores were computed from mKate2$^+$ and SG$^+$ counts as described[23]. To compute LF, double mKate2/Sytox Green positive counts were subtracted from live cell counts.

**AlphaFold modeling.** Protein sequences for StcE and αHER2-eStcE were used as inputs for ColabFold (https://colab.research.google.com/github/sokrypton/ColabFold/blob/main/AlphaFold2.ipynb#scrollTo=kOblAo-xetgx) to obtain model structures[54]. The predicted structure for StcE aligned well (for previously reported StcE structure[31] RMSD=0.2) with the X-ray structure determined by Yu *et al.*[31]. Molecular graphics were generated using PyMOL.

**Cloning.** StcE mutants were cloned from pET28b-StcE_Δ35-NHis, generously provided by Natalie Strynadka (University of British Columbia), using a Q5 Site-Directed Mutagenesis Kit (New England Biolabs), In-Fusion HD Cloning Plus (Takara Bio), or from ordering related designed plasmids from Twist Bioscience. All plasmids were sequence confirmed (Elim Biopharm) before proceeding. The amino acid sequence for the 5F7 nanobody was provided by Melissa Gray[56], reverse translated, and optimized for expression in *Escherichia coli* K12 with the IDT Codon Optimization Tool before cloning as above.

**Protein purification.** BL21(DE3) *Escherichia coli* were transformed with sequence confirmed plasmids and grown in sterile terrific broth with 30 µg/ml kanamycin at 37 °C, 250 rpm until an optical density of 0.4-0.8 was reached. Protein expression was induced with 0.3 mM IPTG and the culture was incubated overnight at 20 °C, 250 rpm. Cells were spun down at 6000 *x g* for 10 min and lysed in 20 mM HEPES, pH 7.5, 500 mM NaCl with a probe tip sonicator. Lysates were clarified by spinning at 11,000 *x g* for 10 min and filtered through a low protein binding 0.22 µm polyethersulfone membrane vacuum filter bottle (Corning). Lysates were applied to 3-4 mL of Ni-NTA agarose (Qiagen) per liter of bacterial culture, washed with 200 mL of 20 mM HEPES, pH 7.5, 500 mM NaCl, 20 mM imidazole, and eluted with 20 mL of 20 mM HEPES, pH 7.5, 500 mM NaCl, 250 mM imidazole per liter of culture. Purified proteins were buffer exchanged into cold PBS either with Zeba Spin Desalting Columns, 7K MWCO, 0.5 mL capacity (Fisher Scientific) or through dialysis with Pierce Slide-A-Lyzer G2 Dialysis Cassettes, 20K MWCO (Fisher Scientific). Protein concentration was determined via NanoDrop and protein purity was determined by SDS-PAGE. Purified protein aliquots were stored at -80 °C and thawed and stored at 4 °C before experiments.

**Endotoxin-free protein purification.** ClearColi BL21(DE3) electrocompetent cells (Lucigen) were transformed with plasmids and grown in sterile LB-Miller culture media with 30 µg/mL kanamycin at 37 °C, 250 rpm until an optical density of 0.4-0.8 was reached. Protein expression was induced with 0.4 mM IPTG and the proteins were purified as above. Proteins were run through Pierce high-capacity endotoxin removal columns (Thermo Fisher Scientific) at least four times following manufacturer's instructions. Endotoxin levels were tested using HEK-Blue™ LPS Detection Kit 2 (InvivoGen) following manufacturer's instructions. All endotoxin levels were confirmed to be below K/M for the maximum dose used *in vivo*, where K is 5 EU/kg and M is the dose of the protein/formulation of interest within a single hour period[57].

***In vitro* mucin cleavage activity assays.** Recombinant C1-INH (Molecular Innovations) was labeled with IRDye 800CW NHS Ester (LI-COR Biosciences) (dye:protein ratio of 0.54) following manufacturer's instructions. Extra dye was removed with Zeba Spin Desalting Columns, 7K MWCO, 0.5 mL capacity (Fisher Scientific). For each reaction, 50 nM of mucinase and 500 nM of recombinant mucin in PBS were combined and incubated at 37 °C for 1 h. 4X NuPAGE LDS Sample Buffer (Fisher Scientific) and dithiothreitol (DTT) (Thermo Fisher Scientific) were added to a final concentration of 1-2X and 1-250 mM, respectively, and boiled at 95 °C for 5 min. Samples were run on a 4 to 12% 18-well Criterion XT Bis-Tris protein gel (Bio-Rad) in XT MOPS (Bio-Rad) at 180 V for 1 h. Gels were imaged using an Odyssey CLx Near-Infrared Fluorescence Imaging System (LI-COR Biosciences). To quantify mucinase activity, product band and C1-INH parent band signal intensities were determined on Image Studio software, and digestion percentage was calculated by dividing the signal from the product bands by the signal from the parent + product bands. For rhMUC16 digestion experiments, recombinant MUC16 (R&D Systems) was left unlabeled but otherwise treated as above. Protein was visualized with AcquaStain Protein Gel Stain (Bulldog-Bio), washed at least three times with ddH$_2$O, and imaged using an Odyssey CLx Near-Infrared Fluorescence Imaging System. For comparison of *in vitro* substrate digestion efficiency of different mutants, 89 µg/mL of unlabeled substrates and 162 nM of mucinase (StcE, StcE$^{W366A}$, ddStcE, eStcE, or αHER2-eStcE) in PBS were combined and incubated overnight at 37 °C. SDS-PAGE gels were run and analyzed as above. Bovine serum albumin (BSA) was purchased from Sigma-Aldrich (A7906-1KG). Fetuin was purchased from Promega (V4961). Recombinantly expressed MUC16, podocalyxin, CD43, and PSGL-1 were purchased from R&D Systems (5609-MU, 1658-PD, 9680-CD, 3345-PS, respectively

***In cellulo* MUC1 cleavage activity assay.** HeLa cells resuspended at 4.5-5.0x$10^5$ cells in 150 µL of complete media were allocated per well to a 96-well ultra-low attachment round bottom plate (Corning). 50 µL of the different StcE mutants in PBS were added to the wells, and the plate was incubated at 37 °C for 1 h. Cells were washed twice with cold FACS buffer containing 2 mM EDTA, once with cold FACS buffer, and stained with anti-MUC1/episialin antibody (clone 214D4) (EMD Millipore) in FACS buffer supplemented with 0.1% benzonase for 30 min on ice. Cells were washed three times with cold FACS buffer containing 2 mM EDTA and stained with Alexa Fluor 647 Affinipure Goat Anti-Mouse IgG (Jackson ImmunoResearch). Cells were washed twice with FACS containing 2 mM EDTA and stained with 30 nM Sytox Green for 10 min at 4 °C prior to analysis using a BD Accuri C6 plus. For data analysis, the Alexa Fluor 647 mean fluorescence intensity of unstained and PBS-treated samples were used to define 0% and 100% cell surface MUC1, respectively, and each sample was normalized to percent MUC1 within each replicate. Using GraphPad Prism 9, each replicate was fitted to inhibitor concentration vs normalized response and $\log_{10}(IC_{50})$ was reported.

**Mucinase binding assays to cell surface mucins.** 4-5x$10^5$ HeLa cells were added to each well of a V-bottom 96-well plate and washed three times with cold FACS buffer containing 2 mM EDTA. Cells were treated with mucinases in cold FACS buffer containing 2 mM EDTA and 0.1% benzonase for 30 min on ice, washed three times with FACS with 2 mM EDTA, and stained with FITC anti-His antibody (clone GG11-8F3.5.1) (Miltenyi Biotec). Cells were washed three times with FACS buffer with 2 mM EDTA and stained with 5 nM Sytox Red in FACS buffer with 2 mM EDTA for 20 min prior to analysis using a MACSQuant Analyzer 10 Flow Cytometer (Miltenyi Biotec). Within each sample, binding was normalized to the greatest mean fluorescence intensity (MFI) per mucinase, with 0 being defined as the MFI for the PBS-treated sample. To mitigate the hook effect, only concentrations with greater than 85% normalized binding past the maximum signal were included. In GraphPad Prism 9, each replicate was fitted to agonist concentration vs response with the lower limit restricted to 0 and $\log_{10}(EC_{50})$ was reported. For conjugate and nanobody binding assays, MCF10A$^{\pm MUC1\ \pm HER2}$ cells were processed, washed, and stained as described above. Cells were analyzed on a BD Accuri C6 plus. For replicates in which Prism could not correctly fit the data to report an EC50 value, the replicate was not included in the bar graph of EC50 values; this occurred with one replicate of αHER2-eStcE binding to MCF10A.

**ddStcE$^{W366A}$ mass spectrometry sample preparation.** Recombinantly expressed MUC16, podocalyxin, CD43, and PSGL-1 were purchased from R&D Systems (5609-MU, 1658-PD, 9680-CD, 3345-PS, respectively). All recombinant mucin-domain glycoproteins were reconstituted in ultrapure water (Pierce) to a concentration of 1 mg/mL. A fraction (1 µg; 1 µL) of each recombinant glycoprotein was digested with ddStcE$^{W366A}$ at a 1:1 enzyme-to-substrate (E:S) ratio. For sialidase-treated samples, 1 µL of sialoEXO (Genovis) was added to 39 µL of ultrapure water, and 1 µL of this dilution was added to the reaction vial, per manufacturer instructions. The reaction was brought to a total volume of 12 µL in 50 mM ammonium bicarbonate and allowed to react overnight at 37 °C. Control reactions were incubated at 37 °C overnight in a solution containing buffer only. The following day, the volume was increased to 19 µL with 50 mM ammonium bicarbonate. PNGaseF (1 µL; Promega) was added to 99 µL of 50 mM ammonium bicarbonate, and 1 µL of this reaction was added to each mucinase reaction vial. De-N-glycosylation reactions were incubated for 8-12 h at 37 °C. Reduction and alkylation were performed according to ProteaseMax (Promega) protocols. Briefly, the solution was diluted to 93.5 µL with 50 mM ammonium bicarbonate. Then, 1 µL of 0.5 M dithiothreitol (DTT) was added and the samples were incubated at 56 °C for 20 min, followed by the addition of 2.7 µL of 0.55 M iodoacetamide at room temperature for 15 min in the dark. Digestion was completed by adding sequencing-grade trypsin (Promega) at a 1:20 E:S ratio overnight at 37 °C and quenched by adding 0.3 µL of glacial acetic acid. C18 clean-up was performed using 1 mL StrataX columns (Phenomenex). Each column was wet with 1 mL of acetonitrile once, followed by one 1 mL rinse of buffer A (0.1% formic acid in water). The samples were diluted to 1 mL in buffer A and loaded through the column, then rinsed with buffer A. Finally, the samples were eluted with three rinses of 100 µL of buffer B (0.5% formic acid, 80% acetonitrile) and dried by SpeedVac. The samples were reconstituted in 10 µL of buffer A for MS analysis.

**Mass spectrometry for cleavage motif.** Samples were analyzed by online nanoflow liquid chromatography-tandem mass spectrometry using an Orbitrap Fusion Tribrid mass spectrometer (Thermo Fisher Scientific) coupled to a Dionex Ultimate 3000 HPLC (Thermo Fisher Scientific). Each sample was analyzed twice; once with an HCD triggered electron transfer dissociation (ETD) method (for input to Byonic), and the second with an HCD triggered EThcD method (for input into OPair). A portion of the sample (4 µL of 10 µL; 40%) was loaded via autosampler isocratically onto a C18 nano precolumn using 0.1% formic acid in water ("solvent A"). For preconcentration and desalting, the column was washed with 2% acetonitrile and 0.1% formic acid in water ("loading pump solvent"). Subsequently, the C18 nano precolumn was switched in line with the C18 nano separation column (75-µm × 250-mm EASYSpray containing 2 µm C18 beads) for gradient elution. The column was held at 40 °C using a column heater in the EASY-Spray ionization source (Thermo Fisher Scientific). The samples were eluted at a constant flow rate of 0.3 µL/min using a 90 min gradient. The gradient profile was as follows (min:% solvent B, 2% formic acid in acetonitrile): 0:3, 3:3, 93:35, 103:42, 104:95, 109:95, 110:3, 140:3. The instrument method used an MS1 resolution of 60,000 full width at half maximum (FWHM) at 400 m/z, an automatic gain control (AGC) target of 3e5, and a mass range from 300 to 1,500 m/z. Dynamic exclusion was enabled with a repeat count of 3, repeat duration of 10 s, and exclusion duration of 10 s. Only charge states 2 to 6 were selected for fragmentation. MS2s were generated at top speed for 3 s. Higher-energy collisional dissociation (HCD) was performed on all selected precursor masses with the following parameters: isolation window of 2 m/z, 30% collision energy, orbitrap detection (resolution of 30,000), and an AGC target of 1e4 ions. For HCD-pd-ET(hc)D runs, ET(hc)D was performed if 1) the precursor mass was between 300 and 1,000 m/z and 2) 3 of 9 HexNAc or NeuAc fingerprint ions (126.055, 138.055, 144.07, 168.065, 186.076, 204.086, 274.092, and 292.103) were present at ± 0.1 m/z and greater than 5% relative intensity. ETD parameters were as follows: calibrated charge-dependent ETD times, 2e5 reagent target, and precursor AGC target 1e4. EThcD parameters were the same but included 30 nce supplemental activation and Orbitrap analysis at a resolution of 30,000 FWHM.

**Mass spectrometry data analysis for cleavage motif.** HCD-pd-ETD raw files were searched using Byonic by ProteinMetrics against directed databases containing the recombinant protein of interest. Search parameters included semi-specific cleavage specificity at the C-terminal site of R and K, meaning non-tryptic cleavage was permitted at either the N- or C-terminus of a detected peptide but not both. Mass tolerance was set at 10 ppm for MS1s, 0.1 m/z for HCD MS2s, 0.35 m/z for ETD MS2s. Methionine oxidation (common 2), asparagine deamidation (common 2), and N-term acetylation (rare 1) were set as variable modifications with a total common max of 3, rare max of 1. O-glycans were also set as variable modifications (common 2), using the "O-glycan 6 most common" database. Cysteine carbaminomethylation was set as a fixed modification. Peptide hits were filtered using a 1% FDR. All peptides were manually validated and/or sequenced using Xcalibur software (Thermo Fisher Scientific). HCD was used to confirm that the peptides were glycosylated and ETD spectra were used for site-localization of glycosylation sites.

To further confirm ddStcE$^{W366A}$ cleavage sites, HCD-pd-EThcD raw files were searched using O-Pair in Metamorpheus against directed databases containing the recombinant protein of interest[58]. Search parameters included an "O-glycopeptide search" using the "Oglycan.gdb" database. The top 50 candidates were kept, using HCD-pd-EThcD fragmentation, with a maximum of 4 glycans allowed. Semi-trypsin cleavage specificity was selected with a maximum of 2 missed cleavages, and a peptide length of 5-60. Mass tolerance was set at 10 ppm for MS1s, and 20 ppm for all MS2s. Cysteine carbaminomethylation was set as a fixed modification, and methionine oxidation and asparagine deamidation were set as variable modifications. O-Pair results were filtered for results with a Q value of less than 0.01.

**TAILS mass spectrometry sample preparation.** TAILS methods were adapted from previous TAILS publications[34,59] and protocols available at the Overall group website, https://clip.ubc.ca/resources/protocols-and-sops/ (Bench Protocol v5.6).

K562$^{HER2}$ were washed three times with warmed PBS, incubated for 1-2 h in serum free RPMI without phenol red and without glutamine, and resuspended in the same media at 0.8 million cells/mL. αHER2-eStcE, eStcE, StcE, or equal volume PBS were added to a final concentration of 1 nM, and the cells were incubated overnight at 37 °C. Cells were spun down at 600xg for 5 min, and conditioned supernatant was collected and treated with protease inhibitors (cOmplete, EDTA-free Protease Inhibitor Cocktail) and 10 mM EDTA. Conditioned supernatant was clarified by centrifugation at 1000xg for 5 min at 4 °C. Trichloroacetic acid was added to a final concentration of 15% (v/v), and the mixture sat on ice for 3-4 h. Precipitated proteins were washed three times by repeated pelleting by centrifugation at 9000xg for 15 min at 4 °C, decanting of the supernatant, and resuspension of the pellets in -20 °C 100% acetone. After the final spin, the supernatant was decanted, and the pellets were frozen overnight.

Pellets were resuspended in 100 uL of 6M guanidine hydrochloride, and protein amounts were determined by BCA protein assay kit (Thermo Fisher Scientific). 125 μg of total protein material was used for each sample, and samples were adjusted to a total volume of 175 μL with water. Samples were then adjusted to 100 mM HEPES before adding freshly prepared TCEP to a final concentration of 10 mM and incubation at 37 °C for 30 min. Freshly prepared N-ethylmaleimide (NEM, adjusted to pH 6) was added to a final concentration of 15 mM, and samples were incubated for 10 min in the dark at room temperature. 50 μL of 100 mM TCEP was added, and samples were vortexed to quench NEM before samples were put on ice. Proteins were precipitated using acetone, where a 4-fold volume of ice-cold acetone was added to each sample in an acetone-compatible tube. Samples were vortexed and incubated at -80 °C overnight before centrifuging at 18,000 x g for 10 min. Supernatant was decanted with care taken not to disturb the protein pellet, and pellets were air dried for 15 min in an uncapped tube. Samples were then labeled with 16-plex Tandem Mass Tags (TMT, Thermo Fisher Scientific) according to the labeling scheme in Supplementary Table 5, with distinct differences from manufacturer protocols because of labeling at protein rather than peptide level. Samples were resuspended in 110 μL 100 mM TEAB, and TMT labels (0.8 mg each) were dissolved in 110 μL DMSO. Samples were vortexed prior to a 1 h incubation in the dark at 25 °C. Following incubation, samples were combined into a single tube, and proteins were precipitated using the same acetone precipitation protocol described above with overnight incubation. Following the 15 min airdry, the pellet was resuspended in 50 μL 6M guanidine hydrochloride and then diluted ten-fold with 100 mM HEPES, pH 8.0. Trypsin was added at a protease:protein ratio of 1:100, with gentle mixing with a pipette prior to overnight incubation at 37 °C. Negative selection for N-terminal peptides was performed using 45 mg/mL HPG-ALDII obtained from the Overall Lab (29 mg aliquot, Lot #002121800521). The HPG-ALDII polymer was thawed at room temperature and added to the digested sample at a polymer:peptide ratio of 6:1. Then sodium cyanoborohydride was immediately added to a final concentration of 20 mM, and the sample was gently mixed, the pH was confirmed to be ~6-7, and the sample was incubated overnight at 37 °C. Sample recovery was performed the following day, with all spins at 12,000 x g for 10 min at room temperature. Tris pH 6.8 was added to a final concentration of 100 mM, pH 6-7 was verified, and the sample was incubated for 30 min at 37 °C. A 10-kDA molecular weight cutoff Amicon column was pre-washed with 400 µL 100 mM NaOH and 400 µL water, and flow-through (FT) was discarded. The peptide-polymer mixture was spun through the column and FT was collected into a clean tube (TAILS sample 1). The filter was then washed by spinning 400 µL water through, this FT was added to TAILS sample 1, and then the filter was thoroughly washed with 100 µL water, which rids the filter of the very hydrophilic polymer. The filter was repositioned upside down in a new tube (TAILS sample 2) with a quick spin to increase the yield of hydrophobic peptides. TAILS sample 1 and sample 2 were lyophilized before they were desalted using 10 mg/1 mL Strata-X columns (Phenomenex). Briefly, columns were wet with 1 mL acetonitrile followed by equilibration with 1 mL 0.2% formic acid (FA) in water. Samples were resuspended in 500 µL 0.2% FA in water and were loaded on the column, followed by a wash with 1 mL 0.2% FA. Peptides were eluted with 400 µL 0.2% FA, 80% acetonitrile, dried via lyophilization, then resuspended in 0.2% FA in water prior to MS analysis.

**Supplementary Table 5. Sample IDs and TMT labeling for TAILS MS.**

| Channel | Sample | Channel | Sample |
|---|---|---|---|
| 126C | PBS 1 | 127N | StcE 1 |
| 127C | PBS 2 | 128N | StcE 2 |
| 128C | PBS 3 | 129N | StcE 3 |
| 129C | PBS 4 | 130N | StcE 4 |
| 130C | eStcE 1 | 131N | αHER2-eStcE 1 |
| 131C | eStcE 2 | 132N | αHER2-eStcE 2 |
| 132C | eStcE 3 | 133N | αHER2-eStcE 3 |
| 133C | eStcE 4 | 134N | αHER2-eStcE 4 |

**TAILS mass spectrometry LC-MS/MS.** Both TAILS sample 1 and sample 2 were analyzed using 90-min LC-MS/MS acquisitions, and TAILS sample 1 was analyzed with an additional 240-min LC-MS/MS acquisition. Peptide mixtures were separated over a 25 cm EasySpray reversed phase LC column (75 µm inner diameter packed with 2 µm, 100 Å, PepMap C18 particles, Thermo Fisher Scientific). The mobile phases (A: water with 0.2% formic acid and B: acetonitrile with 0.2% formic acid) were driven and controlled by a Dionex Ultimate 3000 RPLC nano system (Thermo Fisher Scientific). An integrated loading pump was used to load peptides onto a trap column (Acclaim PepMap 100 C18, 5 um particles, 20 mm length, Thermo Fisher Scientific) at 5 µL/min, which was put in line with the analytical column 5.5 min into the gradient. Gradient elution was performed at 300 nL/min for all analyses. For the 90-min acquisitions, the gradient was held at 0% B for the first 6 min of the analysis, followed by an increase from 0% to 5% B from 6 to 6.5 min, an increase from 5% to 22% B from 6.5 to 66.5 min, an increase from 22% to 90% B from 66.5 to 70 min, isocratic flow at 90% B from 70 to 75 min, and a re-equilibration at 0% B for 15 min. For the 240-min acquisitions, the gradient was held at 0% B for the first 6 min of the analysis, followed by an increase from 0% to 5% B from 6 to 6.5 min, an increase from 5% to 25% B from 6.5 to 200 min, an increase from 25% to 90% B from 200 to 218 min, isocratic flow at 90% B from 218 to 224 min, and a re-equilibration at 0% B for 16 min. For all methods, eluted peptides were analyzed on an Orbitrap Fusion Tribrid MS system (Thermo Fisher Scientific). Precursors were ionized using an EASY-Spray ionization source (Thermo Fisher Scientific) source held at +2.2 kV compared to ground, and the column was held at 40 °C. The inlet capillary temperature was held at 275 °C. Survey scans of peptide precursors were collected in the Orbitrap from 350-1500 Th with an AGC target of 250% (1,000,000 charges), a maximum injection time of 50 ms, and a resolution of 60,000 at 200 m/z. For 90-min analyses, monoisotopic precursor selection was enabled for peptide isotopic distributions, precursors of z = 2-5 were selected for data-dependent MS/MS scans for 2 seconds of cycle time, and dynamic exclusion was set to 30 sec with a ±10 ppm window set around the precursor monoisotope. An isolation window of 1 Th was used to select precursor ions with the quadrupole. MS/MS scans were collected using HCD at 30 normalized collision energy (nce) with an AGC target of 200% (100,000 charges) and a maximum injection time of 118 ms. Mass analysis was performed in the Orbitrap with a resolution of 60,000 with a first mass set at 120 m/z. All sets were the same for 240-min analyses, with the exception of a 3 sec cycle time and a 60 sec dynamic exclusion time.

**TAILS mass spectrometry data analysis.** All raw data files were processed in batch using MaxQuant[60], where the Andromeda search engine[61] was used to search the entire human proteome downloaded from Uniprot (reviewed, 20428 entries). Cleavage specificity was set to “semi-specific free N-terminus” with ArgC specificity. The NEM modification of cysteine had to be created, with an addition of C6H7O2N (125.0478 Da)[62], that was as a fixed modification, while, oxidation methionine was set as a variable modification, with 5 maximum modifications per peptide. The experiment type was set to Reporter ion MS2 with 16-plex TMT modifications selected (user defined modifications added for both Lys and N-terminal labeling). The reporter ion mass tolerance was set to 0.003 Da and the minimum reporter PIF score was set to 0.75. Defaults were used for the remaining settings, including PSM and protein FDR thresholds of 0.01 and 20 ppm, 4.5 ppm, and 20 ppm for first search MS1 tolerance, main search MS1 tolerance, and MS2 product ion tolerance, respectively. “Match between runs” and “second peptide” options were not enabled. Quantified peptides were then processed in Perseus[63]. Contaminants and reverse hits were removed, and signal in all relevant TMT channels of at least one condition was required to retain protein identifications.

The four proteins specifically degraded by StcE as compared to PBS (CD99L2, TNFRSF1B, CD55, CD46) were manually searched for regions with a high density of predicted mucin-type o-glycosylation using NetOGlyc-4.0[13]. To account for phosphorylated residues incorrectly predicted as glycosylated residues, phosphorylation was annotated using PhosphoSitePlus[64].

**Generation of HER2 stable lines.** pMXs-HER2 vector was generated by cloning the HER2+ coding sequence (Addgene) into the pMXs-FLAG backbone using In-Fusion HD Cloning Plus (Takara Bio). 1.5x$10^6$ HEK-293Ts were seeded into 6 cm dishes in 5 mL of complete media. 28 h later, 1 µg of pMXs-HER2 was mixed with 900 ng of retrovirus pol/gag, 150 ng of VSVg DNA, 130 µL of DMEM, and 6 µL of 1 mg/mL polyethylenimine (PEI). The mixture was incubated for 20 min at room temperature and added to HEK-293T cells dropwise. 18 h later, the culture media was replaced with 5 mL of DMEM supplemented with 30% heat inactivated FBS and 1% P/S. 30 h later, the media was collected and spun at 1000 rpm for 5 min. The clarified supernatant was stored at -80 ºC prior to infection. To establish stably expressing cell lines, 1.5x$10^6$ cells were seeded in 6-well plates in 2.8 mL of complete media. Polybrene was added at 10 µg/mL and cells were infected with 200 µL of virus-containing media. Plates were spun at 2200 rpm for 45 min and incubated at 37 ºC, 5% $CO_2$. 12-24 h later, cells were lifted with trypsin and plated in 10 cm dishes with 10 µg/mL blasticidin S (Thermo Fisher Scientific). Stable cell lines were tested for expression of HER2 by flow cytometry.

**HER2 flow cytometry.** Log-phase cells were aliquoted into a V-bottom 96-well plate at 5x$10^5$ cells/well. Cells were washed twice with cold FACS buffer with 2 mM EDTA, once with cold FACS buffer, and stained with Alexa Fluor 488 anti-human CD340 (erbB2/HER-2) antibody

(24D2 clone) in FACS buffer containing 0.1% benzonase on ice protected from light. Cells were washed three times with FACS buffer with 2 mM EDTA and stained with 5 nM Sytox Red for 10 min at 4 °C prior to analysis using a BD Accuri C6 plus.

**K562 mixed cell CD43 cleavage assay.** 2.5x$10^5$ K562 and K562$^{HER2}$ cells were allocated per well to a 96-well ultra-low attachment round bottom plate in 150 µL of complete media. 50 µL of mucinases in PBS were added to wells and the plate was incubated (overnight unless stated otherwise) at 37 °C. Cells were washed twice with cold FACS buffer with 2 mM EDTA, once with cold FACS buffer, and stained with Alexa Fluor 488 anti-human CD340 (erbB2/HER-2) antibody (24D2 clone) (BioLegend) and Alexa Fluor 647 CD43/sialophorin antibody (MEM-59 clone) (Novus Biologicals) in FACS buffer supplemented with 0.1% benzonase on ice protected from light. Cells were washed three times with cold FACS buffer with 2 mM EDTA and stained with 1 µM Sytox AADvanced (Thermo Fisher Scientific) in FACS buffer with 2 mM EDTA for 5 min on ice prior to analysis using a BD Accuri C6 plus. Samples were compensated using single-stained controls in FlowJo v. 10.0. Unstained and PBS-treated samples were used to define 0% and 100% cell surface CD43, respectively, and each sample was normalized to percent CD43 within each replicate. Using GraphPad Prism 9, replicates were fitted to inhibitor concentration vs normalized response.

**MCF10A$^{MUC1}$ mixed cell MUC1 cleavage assay.** MUC1ΔCT was induced with 1 µg/mL doxycycline for 24 h. HER2+ cells were stained with 5 µM Molecular Probes CellTracker Green CMFDA Dye (Thermo Fisher Scientific) in PBS at 37 °C for 30 min and washed twice with warmed PBS. 2.5x$10^5$ MCF10A$^{MUC1, HER2}$ and 2.5x$10^5$ MFC10A$^{MUC1}$ (matched for induction or not with doxycycline) were added to each well of a low adhesion U-bottom 96-well plate in 150 µL of complete media. 50 µL of mucinases in PBS were added to wells and the plate was incubated overnight at 37 °C, 5% $CO_2$. Cells were washed twice with cold FACS buffer with 2 mM EDTA, once with cold FACS buffer, and stained with MUC1 mouse mAb (clone VU4H5) (Cell Signaling Technology) in FACS buffer supplemented with 0.1% benzonase on ice for 30 min. Cells were washed three times with cold FACS buffer with 2 mM EDTA, stained with Alexa Fluor 647 Affinipure Goat Anti-Mouse IgG (Jackson ImmunoResearch) for 30 min on ice, washed three times with cold FACS buffer with 2 mM EDTA, and stained with 1 µM Sytox AADvanced (Thermo Fisher Scientific) in FACS buffer with 2 mM EDTA for 5 min on ice prior to analysis using a BD Accuri C6 plus.

**Macrophage phagocytosis assay.** MCF7 and MCF7$^{HER2}$ cells were lifted with enzyme-free cell dissociation buffer and resuspended in PBS. MCF7 and MCF7$^{HER2}$ cells were incubated in 5 µg/mL Alexa Fluor 546 $C_5$ maleimide (Invitrogen) and 5 µg/mL Alexa Fluor 647 $C_2$ maleimide (Invitrogen), respectively, for 20 min rotating at room temperature. Cells were resuspended in 5 mM N-ethyl-maleimide (Sigma Aldrich) in PBS and incubated for 20 min rotating at room temperature. Cells were resuspended in PBS and treated with PBS, 5 nM endotoxin-free StcE, or 100 nM αHER2-eStcE as appropriate for 2 h at 37 °C. After 1 h, *InVivo*MAb anti-mouse/human/rat CD47 (clone MIAP410) (BioXCell) was added to a final concentration of 20 µg/mL. The media for the macrophages was replaced with serum-free RPMI and appropriate wells were treated with 10 µM cytochalasin D (Invitrogen). MCF7 and MCF7$^{HER2}$ cells were washed twice with PBS and resuspended in 100 µL of serum-free RPMI. Macrophage media was replaced with 200 µL of serum-free RPMI. MCF7 and MCF7$^{HER2}$ cells of the same treatment group were mixed and added to the appropriate macrophage well and incubated for 30 min at 37 °C. After incubation, macrophages were gently washed five times with cold PBS. Cells were fixed with 4% paraformaldehyde in PBS for 15 min at room temperature. Cells were then rinsed with PBS and permeabilized with 0.5% Triton-X-100 in PBS for 10 min. Cells were subsequently rinsed with PBS and blocked with 2% BSA in PBS for 20 min. Cells were then stained with

Alexa Fluor 488 phalloidin (Invitrogen) (1:2000) and 7.5 μM DAPI in PBS for 20 min at room temperature. Cells were then washed three times with PBS and stored in PBS at 4 °C until imaging using a Nikon A1R confocal microscope. Phagocytosis binding indices were calculated as the surface area of target cells divided by the number of macrophages in the field of view. Surface area and number of macrophages were calculated using the imaging software Imaris. Normalized binding indices were calculated relative to the binding index of the PBS treatment condition of the appropriate biological replicate. Three biological replicates were done with macrophages isolated from three different human donors.

***In vivo* toxicity studies.** Experiments involving animals were approved under Stanford APLAC protocol no. 31511 and no. 10266. 10-week-old C57BL/6 mice (bred in-house) were injected with PBS or StcE (0.15 mg/kg or 15 mg/kg) via intravenous injection. 9-week-old female BALB/cJ mice (Jackson Labs) were injected with PBS or 10 mg/kg αHER2-eStcE via intraperitoneal injection. Three h post injection, mice were submitted to the Necropsy Service at Stanford University for necropsy and histopathologic analysis, and blood samples submitted to the Animal Diagnostic Laboratory for complete blood count (CBC) analyses.

***In vivo* biodistribution and mucin degradation Western blots.** Experiments involving animals were approved under Stanford APLAC protocol no. 31511. 12-week-old male BALB/cJ mice (Jackson Labs) were injected with PBS, 10 mg/kg StcE, or 0.25 mg/kg IRdye 800CW-labeled StcE via intraperitoneal injection. Liver, spleen, lung, and plasma (submandibular bleed) were collected at the indicated times post injection. Tissues were lysed using a Bead Mill 24 Homogenizer (Fisher Scientific) in RIPA buffer (Thermo Fisher Scientific) supplemented with benzonase and cOmplete Mini, EDTA-free Protease Inhibitor Cocktail Tablets (Sigma Aldrich). Plasma and tissue lysates (40-50 μg) were loaded onto a 4 to 12% Criterion XT Bis-Tris protein gel and run in XT-MOPS at 180 V for 1 h. Total protein was visualized with AcquaStain protein gel stain (Bulldog-Bio). For mucin Western blots, the gel was transferred to a 0.2-μm nitrocellulose membrane using the Trans-Blot Turbo Transfer System (Bio-Rad) at 2.5 A for 15 min. Total protein was quantified using REVERT stain (LI-COR Biosciences). The membrane was blocked with Carbo-free Blocking Solution (Vector Laboratories) supplemented with 0.1% v/v Tween-20 for 1 h at room temperature and then incubated with 10 μg/mL biotin-StcE$^{E447D}$ in PBS-T (0.1% v/v Tween-20) at room temperature for 1 h. IRDye 800CW streptavidin (LI-COR Biosciences) was used according to manufacturer recommendations. Blots were imaged using an Odyssey CLx Near-Infrared Fluorescence Imaging System (LI-COR Biosciences).

8-week-old female BALB/cJ mice (Jackson Labs) were injected with PBS or IRdye 680RD-αHER2-eStcE at 0.25, 0.5, 1, 2, 5, and 10 mg/kg via retro-orbital injection. Plasma (tail bleed) was collected at t=1, 3, 6, 20, and 48 h post injection. Liver, kidney, spleen, lung, and heart tissues for the mice injected with 10 mg/kg αHER2-eStcE were collected at t=20 and 48 h post injection. Plasma samples (1 μL) were loaded onto a 4 to 12% Criterion XT Bis-Tris protein gel and run in XT-MOPS at 180 V for 1 h. Tissues were lysed as described above and lysates (30 μg) were loaded onto a 4 to 12% Criterion XT Bis-Tris protein gel and run in XT-MOPS at 180 V for 1 h. Total protein was visualized with AcquaStain protein gel stain (Bulldog-Bio). Gels were imaged using an Odyssey CLx Near-Infrared Fluorescence Imaging System. 7-week-old female BALB/cJ mice (Jackson Labs) were injected with PBS, 5 mg/kg StcE, or 5 mg/kg αHER2-eStcE via retro-orbital injection. Liver, spleen, lung, and plasma (submandibular bleed) were collected 4 h post injection. Mucin Western blot was performed as described above.

**4T07$^{MUC1, HER2}$ mouse model.** Experiments involving animals were approved under UCSF Institutional Animal Care and Use Committee (IACUC) protocol no. AN179766. 4T07 cells expressing a cytoplasmic truncation of MUC1 (MUC1ΔCT, also referred to as *MUC1* *ectodomain*) were used to limit any possible cytoplasmic signaling[20]. 1x10$^6$ 4T07$^{HER2}$ breast cancer cells expressing mApple luciferase and doxycycline-inducible MUC1ΔCT were seeded in the lungs of female, syngeneic 8-week-old BALB/cJ mice by intravenous injection (tail vein). Cells were stimulated with 2 mg/mL doxycycline for 24 h to induce MUC1ΔCT expression prior to injection. Competent cell seeding of the lungs was assessed by bioluminescent imaging (BLI; IVIS In Vivo Imaging System) following intraperitoneal injection of D-Luciferin (150 mg/kg) within 30 min of cell injection. Mice were given systemic treatments of PBS or 10 mg/kg αHER2-eStcE by intravenous injection every 2 days for a total of 7 doses. Tumor cell lung burden was assessed over time by additional BLI measurements of mice. BLI for independent images was calculated from total bioluminescence flux for the chest region of each mouse. At 15 days, animals were sacrificed, and lungs were harvested. Whole animal and gross lung weights were recorded, as well as the number of detected lung surface lesions. Lungs were then formalin fixed and processed for paraffin embedding and the average lung lesion area for each animal was determined from H&E-stained tissue sections.

**EMT6$^{HER2}$ mouse model.** BALB/c mice were obtained from Janvier Laboratories and bred in-house at the University Hospital Basel, Switzerland. All mouse experiments were approved by the local ethics committee (Approval 2370, Basel Stadt, Switzerland). Animals were housed under specific pathogen-free conditions. For tumor growth experiments, 8-12-week-old females were used. 1x10$^6$ EMT6$^{HER2}$ cells were injected into the right mammary fat pad of female BALB/c mice. For efficacy studies, four I.P. doses of PBS, 10 mg/kg αHER2-eStcE, or an equimolar quantity (2.8 nmol) of αHER2 were administered every 2 days for a total of 4 doses once the tumor size reached an average size of 80-100 mm$^3$. For analysis of the tumor infiltration by flow cytometry, two I.P. doses of PBS, 10 mg/kg αHER2-eStcE or αHER2 were administered every 2 days for a total of 2 doses once the tumor size reached an average size of 80-100 mm$^3$. Perpendicular tumor diameters were measured by caliper and tumor volume calculated according to the following formula: tumor volume (mm$^3$) = ($d^2 \times D$)/2, where $d$ and $D$ are the shortest and longest diameters of the tumor (in millimeters), respectively. Mice were euthanized once tumor size reached approximately 1500 mm$^3$ or when the mice developed ulcerated tumors that required euthanasia and the animals excluded from further analysis.

**Immunohistochemistry of lung tissues for the 4T07$^{MUC1, HER2}$ mouse model.** IHC was performed as previously described[65] using antibodies specific to phospho-FAK tyrosine 397 and Cyclin D1 (see Supplementary Table 3). Briefly, antigen retrieval was accomplished by boiling sections in 10 mM citrate buffer (10min). Following primary antibody incubation overnight at 4 °C, sections were incubated for 1 hr with species-specific Horseradish Peroxidase (HRP)-conjugated secondary antibodies (ImmPRESS HRP Goat Anti-Rabbit IgG Polymer Detection Kit, Peroxidase, Vector Laboratories, Cat. #: MP-7452 and MP-7451) before developing positive staining with ImmPACT DAB Substrate Peroxidase (HRP, Vector Laboratories, Cat. #: SK-4105). Images of stained sections were acquired with an Olympus microscope (IX81) 10x objective and 1.5x magnification.

**Quantitative histological analysis of lung tissues for the 4T07$^{MUC1, HER2}$ mouse model.** Analysis of IHC and H&E-stained lung tissue sections was performed using ImageJ and QuPath software, respectively. Specifically, an IHC profiler ImageJ plugin was used to quantify percent-positive DAB-staining area of lung metastases with selection for either cytoplasmic (phospho-FAK) or nuclear (Cyclin D1) staining[66]. Reported values correspond to the sum of high positive and positive DAB signal. To assess metastatic lesion area throughout the depth of mouse lung tissues, lungs were sectioned as two steps spaced 40 μm apart, with 15 sequential sections of 5 μm cut at each step. The top and bottom sections from each step were then stained with H&E

(75μm apart) and slides were scanned using a ZEISS Axio Scan.Z1 digital slide scanner equipped with CMOS and color cameras and 10x, 20x and 40x objectives. Percent area of lung metastasis for each section was determined in Qupath using the polygon tool to trace and annotate lung lesion area compared to whole tissue section area. The average values of two lung sections from each animal was presented.

**Analysis of *in vivo* mucin cleavage in EMT6$^{HER2}$ mouse model.** Single cell suspensions processed and frozen above were briefly thawed in a 37 °C water bath and immediately placed on ice. Cells were washed once, counted, and 1.9-3x$10^6$ cells were processed per biological sample. Cells were washed once with cold FACS buffer, treated with Mouse BD Fc Block in cold FACS buffer for 5 min on ice, and immediately stained with Brilliant Violet 421 CD45 antibody (30-F11), Alexa Fluor 488 HER2 antibody (24D2), and Alexa Fluor 647 StcE$^{E447D}$ (5 µg/mL) in FACS buffer with 1:1000 benzonase for 30 min on ice protected from light. UltraComp eBeads Plus Compensation Beads (Thermo Fisher Scientific) were stained in parallel for antibody single color controls following manufacturer's recommendations. Cells were washed once with cold PBS and stained with 1:1000 GloCell Fixable Viability Dye Violet 510 (StemCell Technologies) in PBS for 30 min on ice protected from light. ArC Amine Reactive Compensation Beads (Thermo Fisher Scientific) were stained in parallel for viability single color controls following manufacturer's recommendations. Cells were washed once with cold FACS buffer with 2 mM EDTA, resuspended in cold FACS buffer with 2 mM EDTA, and analyzed using MACSQuant Analyzer 10 Flow Cytometer (Miltenyi Biotec). EMT6$^{HER2}$ and immune cells were gated from live single cells as shown in Fig. S12e using fluorescence minus one controls. Correct gating of these populations was confirmed with EMT6$^{HER2}$ and commercial mouse PBMCS (IQ Biosciences) stained and analyzed in parallel with the tumor samples.

**Flow cytometry analysis of tumor infiltrating immune cells**. For the preparation of single cell suspensions, tumors were collected, surgical specimens were mechanically dissociated and subsequently digested using Accutase (PAA Laboratories), collagenase IV (Worthington), hyaluronidase (Sigma) and DNase type IV (Sigma) for 1 h at 37 °C under constant agitation. Cell suspensions were filtered through a 70 µm mesh, 10 μl of CountBright™ Plus absolute counting beads (Invitrogen) were added, and samples were frozen (-80 °C) for further analysis of tumor-infiltrating immune cells by flow cytometry. Thawed samples were stained with antibodies shown above in Supplementary Table 4 and analyzed on Cytek® Aurora instrument. Live single cells were gated for different immune subsets as diagrammed in Fig. S14a. For tSNE analysis, live single CD45+ cells were randomly down sampled using the FlowJo DownSample v3.3.1 plugin. tSNE analysis was performed using the FlowJo tSNE plugin once on the concatenated files using all compensated fluorophores (default settings: learning configuration = opt-SNE, iterations = 1000, perplexity = 30, learning rate = 6996, KNN algorithm = exact(vantage point tree), gradient algorithm = Barnes-Hut) on ~100,000 total cells evenly disturbed between treatment groups and evenly distributed between biological replicates in each treatment group. For analysis of activation states, gates for positive staining were defined using unstained samples and were kept consistent for all immune subsets (Fig. S14b).

### Supplemental Figures and Figure Captions

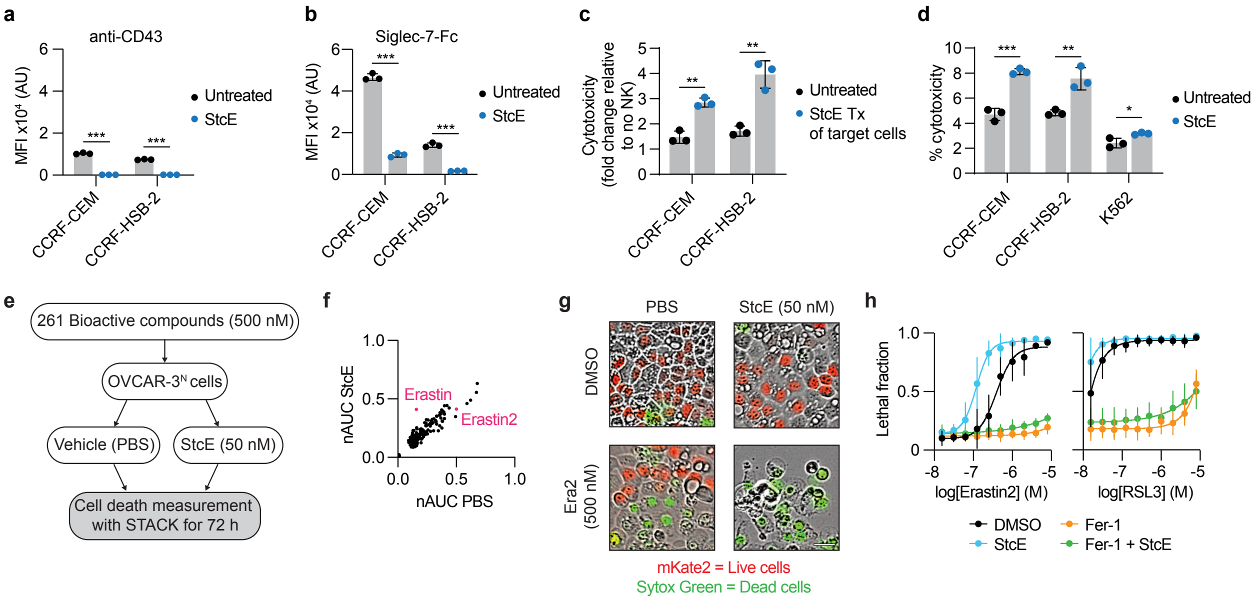

**Fig. S1. StcE treatment of cell lines potentiates NK cell surveillance and small molecule-induced ferroptosis.**

**a-b,** Surface CD43 (**a**) and Siglec-7 ligand (**b**) levels of CCRF-CEM and CCRF-HSB-2 cells ± 50 nM StcE measured by flow cytometry (*n*=3 biological replicates). Siglec-7-Fc staining of K562 cells can be found in Wisnovsky *et al.* (2021).[5]

**c,** Normalized NK cell killing of CCRF-CEM and CCRF-HSB-2 cells ± 10 nM StcE at 5:1 and 2:1 effector:target ratios, respectively (*n*=3 biological replicates). Tx = treatment.

**d,** Cytotoxicity of StcE treatment of leukemia cell lines (*n*=3 biological replicates). A statistically significant baseline toxicity increase was observed.

**e,** Strategy screening for bioactive compound library on OVCAR-3$^N$ cells ± StcE. Superscript *N* denotes stable expression of nuclear fluorescent protein.

**f,** Normalized area-under-the-curve (nAUC) of lethal fraction scores of OVCAR-3$^N$ cells treated with 500 nM bioactive compounds ± 50 nM StcE. Ferroptosis-inducing erastin and erastin2 are highlighted in pink.

**g,** Visualization of live (red) and dead (green) OVCAR-3$^N$ cells ± 50 nM StcE ± 500 nM erastin2 (Era2) at 72 h. Scale bar, 30 μm.

**h,** Lethal fraction curves of OVCAR-3$^N$ cells treated with 500 nM erastin2 or RSL3 ± 50 nM StcE ± 1 μM ferrostatin-1 (Fer-1) for 48 h (*n*=5 biological replicates).

Data are mean ± s.d. *P*-values were determined using Tukey-corrected two-way ANOVA. **p* < 0.05, ***p* < 0.005, ****p* < 0.0005.

**a**

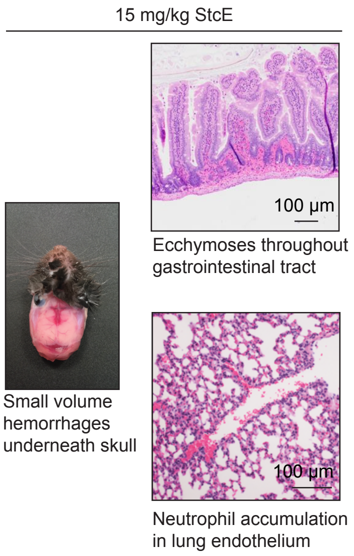


**b**

I.V. injection, 3 h, C57BL/6, *n*=1

| | *Control* | *0.15 mg/kg StcE* | *15 mg/kg StcE* | *Reference Ranges* |
|---|---|---|---|---|
| *White blood cell (WBC)* | 5.86 | 4.44 | 3.94 | 5.5 - 9.3 K/µL |
| *Red blood cell (RBC)* | 10.85 | 9.78 | 13.11 | 7.7 - 8.8 M/µL |
| *Hemoglobin (HGB)* | 15.8 | 14 | 18.3 | 13.7 - 16.4 gm/dL |
| *Hematocrit (HCT)* | 50.3 | 45.5 | 58.9 | 39.0 - 47.0% |
| *Mean corpuscular volume (MCV)* | 46.4 | 46.5 | 44.9 | 52.0 - 68.7 fL |
| *Mean corpuscular hemoglobin (MCH)* | 14.6 | 14.3 | 14 | 18.4 - 19.6 pg |
| *MHC concentration (MCHC)* | 31.4 | 30.8 | 31.1 | 34.0 - 36.0 g/dL |
| *Platelet count* | 1241 | 1345 | 82 | 675 - 1338 K/µL |
| *Red cell distribution width (RDW)* | 20 | 19.2 | 22.2 | % |
| *Platelet distribution width (PDW)* | 7.8 | 8.9 | 7.2 | fL |
| *Mean platelet volume (MPV)* | 6.6 | 7.3 | 6.8 | fL |
| *Platelet-large cell ratio (P-LCR)* | 4.9 | 8.7 | 5.5 | % |
| *Procalcitonin (PCT)* | 0.82 | 0.98 | 0.06 | % |
| *Reticulocyte count* | 4.88 | 4.52 | 6.56 | 1.0 - 2.8% |
| *Immature reticulocyte fraction (IRF)* | 50.3 | 54.9 | 52.3 | % |
| *Low fluorescence reticulocytes (LFR)* | 49.7 | 45.1 | 47.7 | % |
| *Medium fluorescence reticulocytes (LFR)* | 24.6 | 18.7 | 28 | % |
| *Reticulocyte absolute* | 529480 | 442056 | 860016 | /µL |
| *Platelet estimate* | Adequate | Adequate | Decreased | K/µL |
| *Neutrophils* | 4 | 4 | 54 | 15 - 32% |
| *Lymphocytes* | 93 | 92 | 37 | 65 - 83% |
| *Monocytes* | 2 | 3 | 8 | 0 - 3% |
| *Eosinophils* | 1 | 1 | 1 | 0 - 3% |
| *Basophils* | 0 | 0 | 0 | % |
| *RBC morphology* | Normal | Normal | Normal | |
| *Neutrophil absolute* | 234 | 178 | 2128 | 825 - 2604 |
| *Lymphocyte absolute* | 5450 | 4085 | 1458 | 3685 - 7812 |
| *Monocyte absolute* | 117 | 133 | 315 | 0 - 279 |
| *Eosinophil absolute* | 59 | 44 | 39 | 0 - 279 |
| *Basophil absolute* | 0 | 0 | 0 | N/A |

**c**

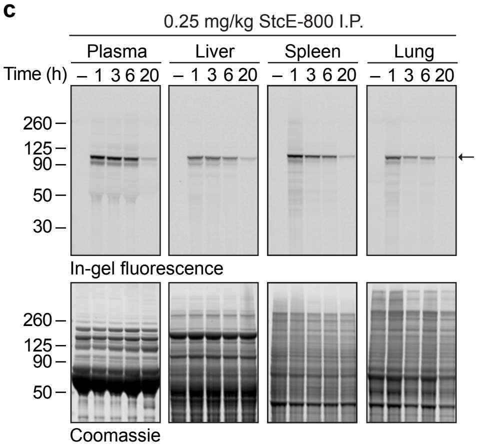


**d**

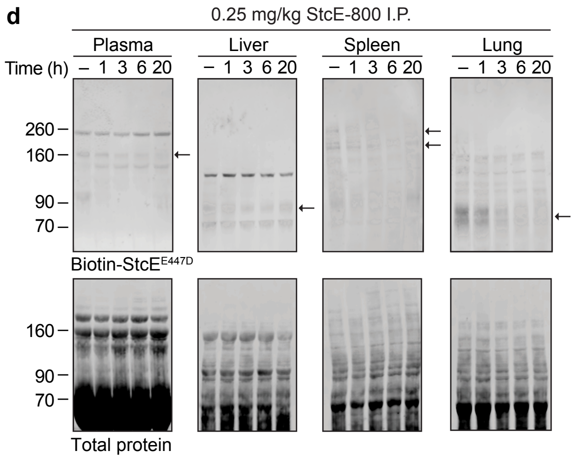


**e**

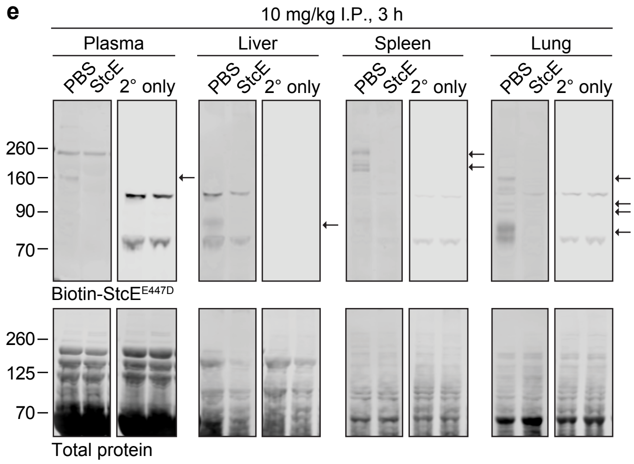

**Fig. S2. StcE cleaves mucins in mouse tissues at a maximum tolerated dose of 0.25 mg/kg and exhibits systemic toxicity at higher doses.**
**a,** Necropsy analysis post intravenous (I.V.) injection of 15 mg/kg StcE revealed abnormalities in the lung, gastrointestinal tract, and underneath the skull (*n*=1).
**b,** Complete blood count (CBC) analyses post I.V. injection of vehicle control (PBS), StcE at 0.15 mg/kg, or StcE at 15 mg/kg (*n*=1).
**c,** SDS-PAGE of plasma and tissues post intraperitoneal (I.P.) injection of PBS or 0.25 mg/kg of IRdye 800CW-labeled StcE (StcE-800, molecular weight = 98 kDa), indicated by the black arrow.
**d,** Mucin Western blot on plasma and tissues from (**c**). Mucin bands are indicated by black arrows.
**e,** Mucin Western blot on plasma and tissues 3 h post I.P. injection of PBS or 10 mg/kg StcE. Mucin bands are denoted by black arrows.

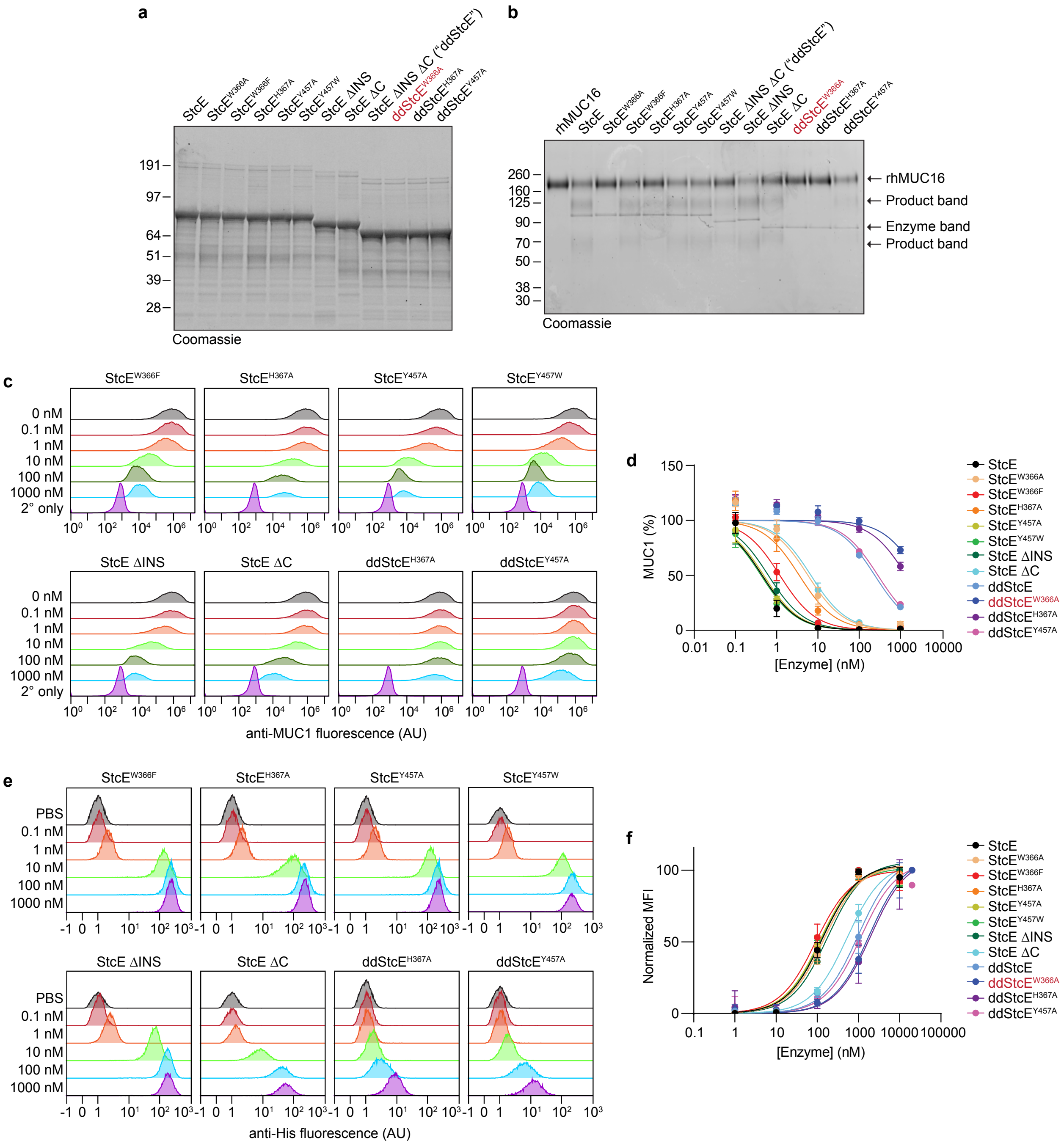


**Fig. S3. Expression and characterization of engineered StcE mutants.**

**a,** SDS-PAGE of purified StcE and StcE mutants.

**b,** Digestion of recombinant MUC16 (rhMUC16) with 50 nM StcE or StcE mutants.

**c,** Representative flow plots related to Fig. 2e-f showing surface MUC1 levels of HeLa cells treated with StcE variants at indicated concentrations.

**d,** MUC1 cleavage curves for StcE and StcE mutants corresponding to Fig. 2e-f and (**c**) (*n*=3 biological replicates).

**e,** Representative flow plots related to Fig. 2g-h depicting cell surface binding of StcE variants on HeLa cells measured by anti-His staining (*n*=3 biological replicates).

**f,** Binding curves for StcE and StcE mutants corresponding to Fig. 2g-h and (**e**) (*n*=3 biological replicates).
Data are mean ± s.d.

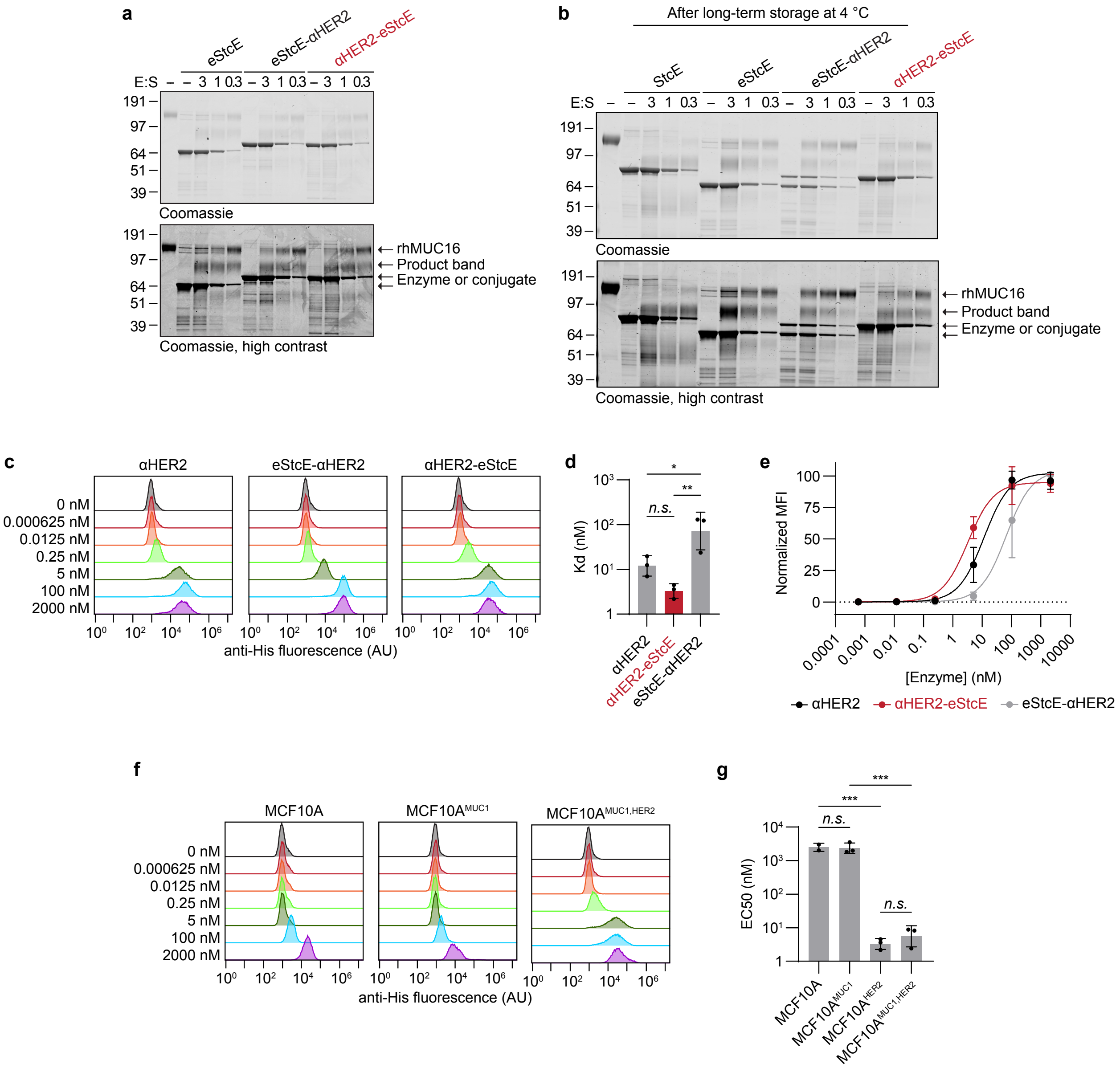


**Fig. S4. Expression and characterization of engineered nanobody-mucinase conjugates.**
**a,** Digestion of rhMUC16 with eStcE alone or nanobody-eStcE conjugates.
**b,** rhMUC16 in-gel digest depicting degradation of eStcE-*αHER2* conjugate after long-term storage at 4 ºC.
**c,** Representative flow plots showing cell surface binding of nanobody alone and eStcE-*αHER2* on MCF10A$^{HER2}$ cells measured by anti-His staining.
**d,** Kd values derived from (**c**) (*n*=3 biological replicates).
**e,** Cell surface binding curves derived from (**c**) (*n*=3 biological replicates).
**f,** Representative flow plots showing cell surface binding of *αHER2*-eStcE on MCF10A$^{\pm MUC1, \pm HER2}$ cells measured by anti-His staining. For flow plot of *αHER2-eStcE on* MCF10A$^{HER2}$, *see Fig. S4c.*
**g,** Kd values derived from (**f**) (*n*=3 biological replicates).
Data are mean ± s.d.

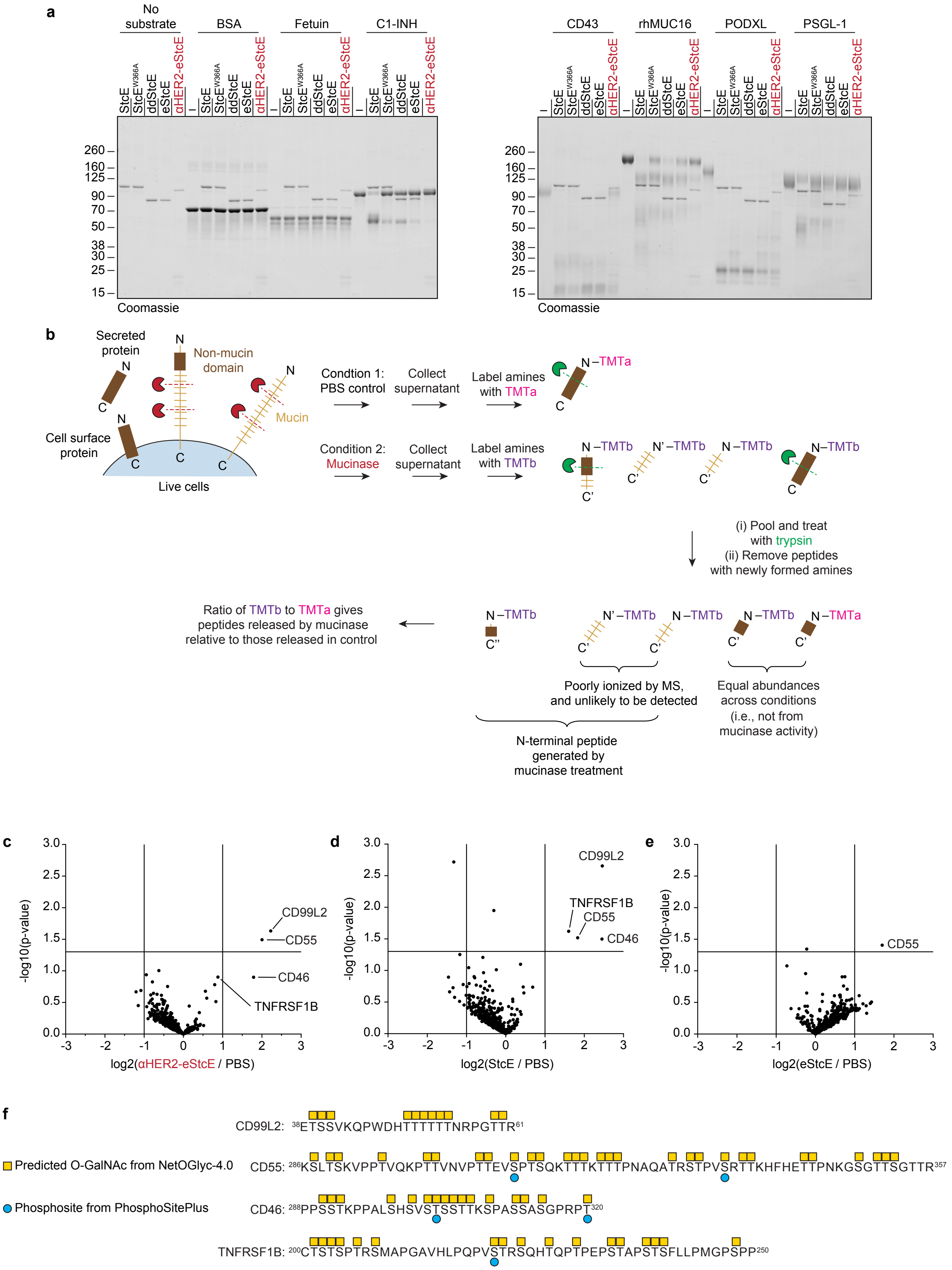
a
No substrate
BSA
Fetuin
C1-INH
CD43
rhMUC16
PODXL
PSGL-1
StcE
StcE W366A
ddStcE
eStcE
αHER2-eStcE
260
160
125
90
70
50
38
30
25
15
Coomassie
b
Secreted protein
Non-mucin domain
Mucin
Cell surface protein
Live cells
Condtion 1: PBS control
Collect supernatant
Label amines with TMTa
Condition 2: Mucinase
Collect supernatant
Label amines with TMTb
N –TMTa
N –TMTb
N' –TMTb
(i) Pool and treat with trypsin
(ii) Remove peptides with newly formed amines
Ratio of TMTb to TMTa gives peptides released by mucinase relative to those released in control
Poorly ionized by MS, and unlikely to be detected
Equal abundances across conditions (i.e., not from mucinase activity)
N-terminal peptide generated by mucinase treatment
c
d
e
-log10(p-value)
log2(αHER2-eStcE / PBS)
log2(StcE / PBS)
log2(eStcE / PBS)
CD99L2
CD55
CD46
TNFRSF1B
f
Predicted O-GalNAc from NetOGlyc-4.0
Phosphosite from PhosphoSitePlus
CD99L2: 38ETSSVKQPWDHTTTTTTNRPGTTR61
CD55: 286KSLTSKVPPTVQKPTTVNVPTTEVSPTSQKTTTKTTTPNAQATRSTPVSRTTKHFHETTPNKGSGTTSGTTR357
CD46: 288PPSSTKPPALSHSVSTSSTTKSPASSASGPRPT320
TNFRSF1B: 200CTSTSPTRSMAPGAVHLPQPVSTRSQHTQPTPEPSTAPSTSFLLPMGPSPP250

**Fig. S5. Assessment of *αHER2*-eStcE selectivity for mucin substrates *in vitro* and on cell surfaces.**
**a,** Digestion of recombinant or purified non-mucins (BSA, fetuin) and mucins (C1-INH, CD43, PODXL, PSGL-1) with StcE, StcE mutants, and *αHER2*-eStcE.
**b,** Setup for terminal amine isotopic labeling of substrates mass spectrometry (TAILS MS) experiment. Mucinase-generated peptides derived from mucin domains were not searched for because of search space complications caused by glycan modifications.
**c-e,** Volcano plots depicting enrichment of peptides following treatment of K562$^{HER2}$ cells with *αHER2*-eStcE (**c**), StcE (**d**), or eStcE (**e**) relative to vehicle control (*n*=4 biological replicates).
**f**, Annotation of predicted O-glycosites (yellow squares)[13] and known phosphosites (blue circles)[64] in putative mucin domains of enriched proteins from (**c-e**).

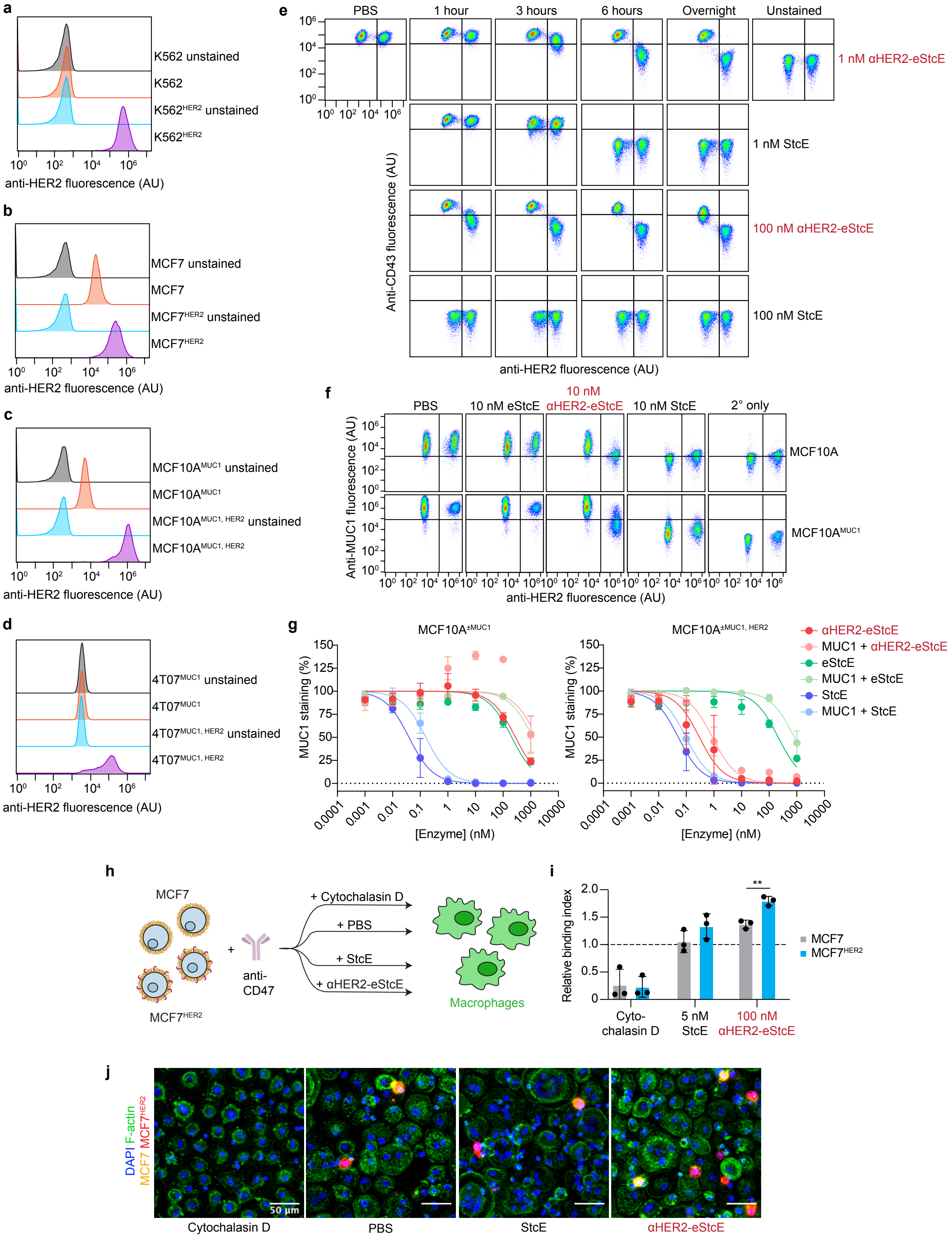
a
K562 unstained
K562
K562HER2 unstained
K562HER2
anti-HER2 fluorescence (AU)
b
MCF7 unstained
MCF7
MCF7HER2 unstained
MCF7HER2
anti-HER2 fluorescence (AU)
c
MCF10AMUC1 unstained
MCF10AMUC1
MCF10AMUC1, HER2 unstained
MCF10AMUC1, HER2
anti-HER2 fluorescence (AU)
d
4T07MUC1 unstained
4T07MUC1
4T07MUC1, HER2 unstained
4T07MUC1, HER2
anti-HER2 fluorescence (AU)
e
PBS
1 hour
3 hours
6 hours
Overnight
Unstained
1 nM αHER2-eStcE
1 nM StcE
100 nM αHER2-eStcE
100 nM StcE
Anti-CD43 fluorescence (AU)
anti-HER2 fluorescence (AU)
f
PBS
10 nM eStcE
10 nM αHER2-eStcE
10 nM StcE
2° only
MCF10A
MCF10AMUC1
Anti-MUC1 fluorescence (AU)
anti-HER2 fluorescence (AU)
g
MCF10A±MUC1
MCF10A±MUC1, HER2
MUC1 staining (%)
[Enzyme] (nM)
αHER2-eStcE
MUC1 + αHER2-eStcE
eStcE
MUC1 + eStcE
StcE
MUC1 + StcE
h
MCF7
MCF7HER2
anti-CD47
+ Cytochalasin D
+ PBS
+ StcE
+ αHER2-eStcE
Macrophages
i
Relative binding index
**
MCF7
MCF7HER2
Cyto-chalasin D
5 nM StcE
100 nM αHER2-eStcE
j
DAPI F-actin MCF7 MCF7HER2
50 µm
Cytochalasin D
PBS
StcE
αHER2-eStcE

**Fig. S6. Generation of HER2+ cell lines and mixed cell assays to assess targeted de-mucination.**

**a-d,** Surface HER2 levels of K562$^{\pm HER2}$ (**a**), MCF7$^{\pm HER2}$ (**b**), MCF10A$^{MUC1, \pm HER2}$ (**c**), and 4T07$^{MUC1, \pm HER2}$ (**d**) cells measured by flow cytometry.

**e,** Representative flow plots depicting surface CD43 levels of mixed K562$^{\pm HER2}$ cells treated with StcE or conjugate for the indicated times and concentrations.

**f,** Representative flow plots depicting surface MUC1 levels of mixed MCF10A$^{\pm MUC1, \pm HER2}$ cells treated with 10 nM mucinases or conjugate.

**g,** MUC1 cleavage curves derived from (**f**) (*n*=3 biological replicates).

**h,** Setup for mixed cell macrophage phagocytosis assay using MCF7$^{\pm HER2}$ cells.

**i,** Relative binding index of MCF7$^{\pm HER2}$ cells treated with cytochalasin D, StcE, or *αHER2*-eStcE (*n*=3 biological replicates).

**j,** Representative confocal microscopy images used for (**i**).

Data are mean ± s.d. *P*-values were determined using t-tests. *$p < 0.05$, **$p < 0.005$, ***$p < 0.0005$.

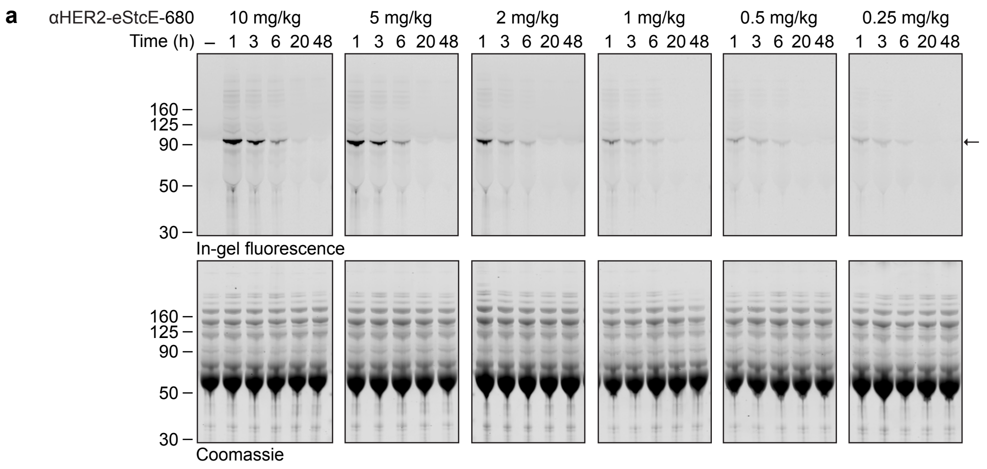


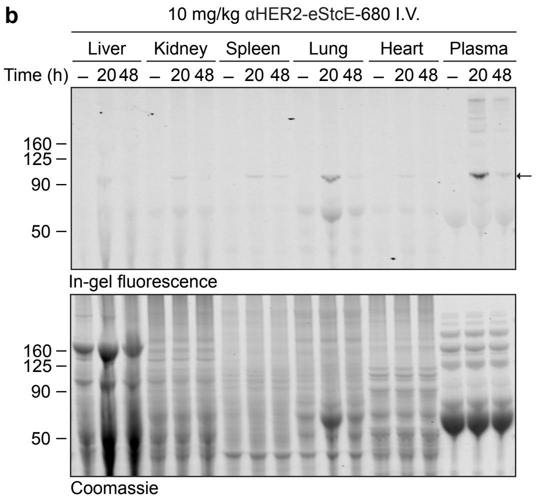


**c**

I.P. injection, 3 h, BALB/c, average of *n*=2

| | *Control* | *10 mg/kg αHER2-eStcE* | *Reference Ranges* |
|---|---|---|---|
| *White blood cell (WBC)* | 5.55 | 5.60 | 5.5 - 9.3 K/μL |
| *Red blood cell (RBC)* | 11.23 | 10.42 | 7.7 - 8.8 M/μL |
| *Hemoglobin (HGB)* | 17.3 | 15.95 | 13.7 - 16.4 gm/dL |
| *Hematocrit (HCT)* | 51.85 | 48.20 | 39.0 - 47.0% |
| *Mean corpuscular volume (MCV)* | 46.20 | 46.25 | 52.0 - 68.7 fL |
| *Mean corpuscular hemoglobin (MCH)* | 15.40 | 15.30 | 18.4 - 19.6 pg |
| *MHC concentration (MCHC)* | 33.35 | 33.10 | 34.0 - 36.0 g/dL |
| *Platelet count* | 1121 | 1030 | 675 - 1338 K/μL |
| *Red cell distribution width (RDW)* | 21.45 | 20.45 | % |
| *Platelet distribution width (PDW)* | 13.20 | 13.40 | fL |
| *Mean platelet volume (MPV)* | 8.50 | 8.60 | fL |
| *Platelet-large cell ratio (P-LCR)* | 19.85 | 20.15 | % |
| *Procalcitonin (PCT)* | 0.96 | 0.88 | % |
| *Reticulocyte count* | 4.16 | 3.95 | 1.0 - 2.8% |
| *Immature reticulocyte fraction (IRF)* | 46.25 | 47.90 | % |
| *Low fluorescence reticulocytes (LFR)* | 53.75 | 52.10 | % |
| *Medium fluorescence reticulocytes (LFR)* | 16.95 | 17.15 | % |
| *Reticulocyte absolute* | 466118 | 410464 | /μL |
| *Platelet estimate* | Adequate | Adequate | K/μL |
| *Neutrophils* | 14 | 33 | 15 - 32% |
| *Lymphocytes* | 82 | 54 | 65 - 83% |
| *Monocytes* | 3.5 | 13.5 | 0 - 3% |
| *Eosinophils* | 1 | 0 | 0 - 3% |
| *Basophils* | 0 | 0 | % |
| *RBC morphology* | Normal | Normal | |
| *Neutrophil absolute* | 767 | 1835 | 825 - 2604 |
| *Lymphocyte absolute* | 4533 | 3006 | 3685 - 7812 |
| *Monocyte absolute* | 197 | 760 | 0 - 279 |
| *Eosinophil absolute* | 59 | 0 | 0 - 279 |
| *Basophil absolute* | 0 | 0 | N/A |

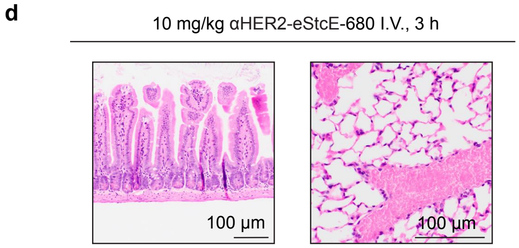


No signs of previously observed gut (left) and lung (right) toxicity

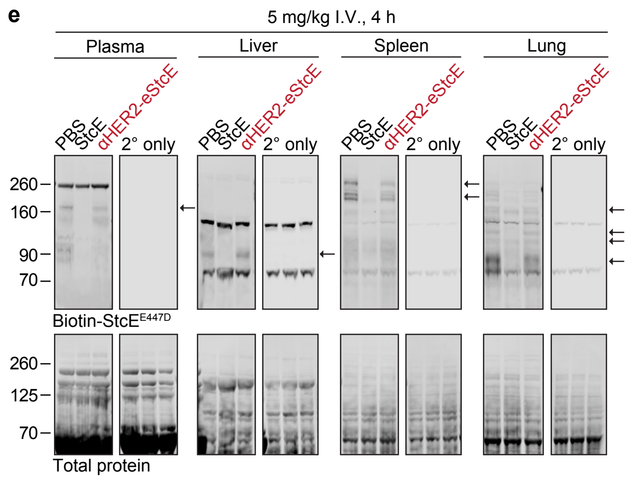

**Fig. S7. *αHER2*-eStcE is nontoxic to mice at every tested dose and distributes widely across tissues.**
**a,** SDS-PAGE of plasma from mice post retro-orbital injection of PBS or IRdye 680RD-labeled *αHER2*-eStcE (*αHER2*-eStcE-680) at the indicated doses. *αHER2*-eStcE is indicated by the black arrow.
**b,** SDS-PAGE of plasma and tissues post retro-orbital injection of 10 mg/kg *αHER2*-eStcE-680, indicated by the black arrow.
**c,** Complete blood count analyses 3 hours post I.P. injection of vehicle control (PBS) or *αHER2*-eStcE at 10 mg/kg (average of *n*=2 biological replicates).
**d,** Necropsy analysis 3 h post retro-orbital injection of 10 mg/kg *αHER2*-eStcE-680 revealed no abnormalities.
**e,** Mucin Western blot on plasma and tissues 4 hours post retro-orbital injection of 5 mg/kg StcE or conjugate. Mucin bands are denoted by black arrows.

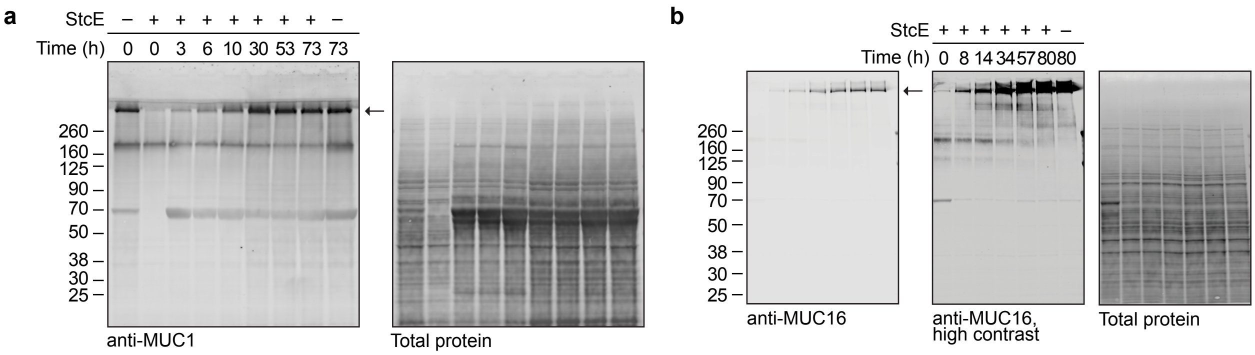


**Fig. S8. Mucin ectodomains cleaved by StcE return within 24 hours.**
**a-b**, 4T07$^{MUC1}$ cells (**a**) and OVCAR-3 cells (**b**) were treated with 50 nM StcE for 2 hours, washed 1x with 2 mM EDTA followed by 5x with DPBS, then cultured for the indicated times. Cells were then lysed and subjected to Western blotting for MUC1 (**a**) and MUC16 (**b**). Mucin bands are denoted by black arrows.

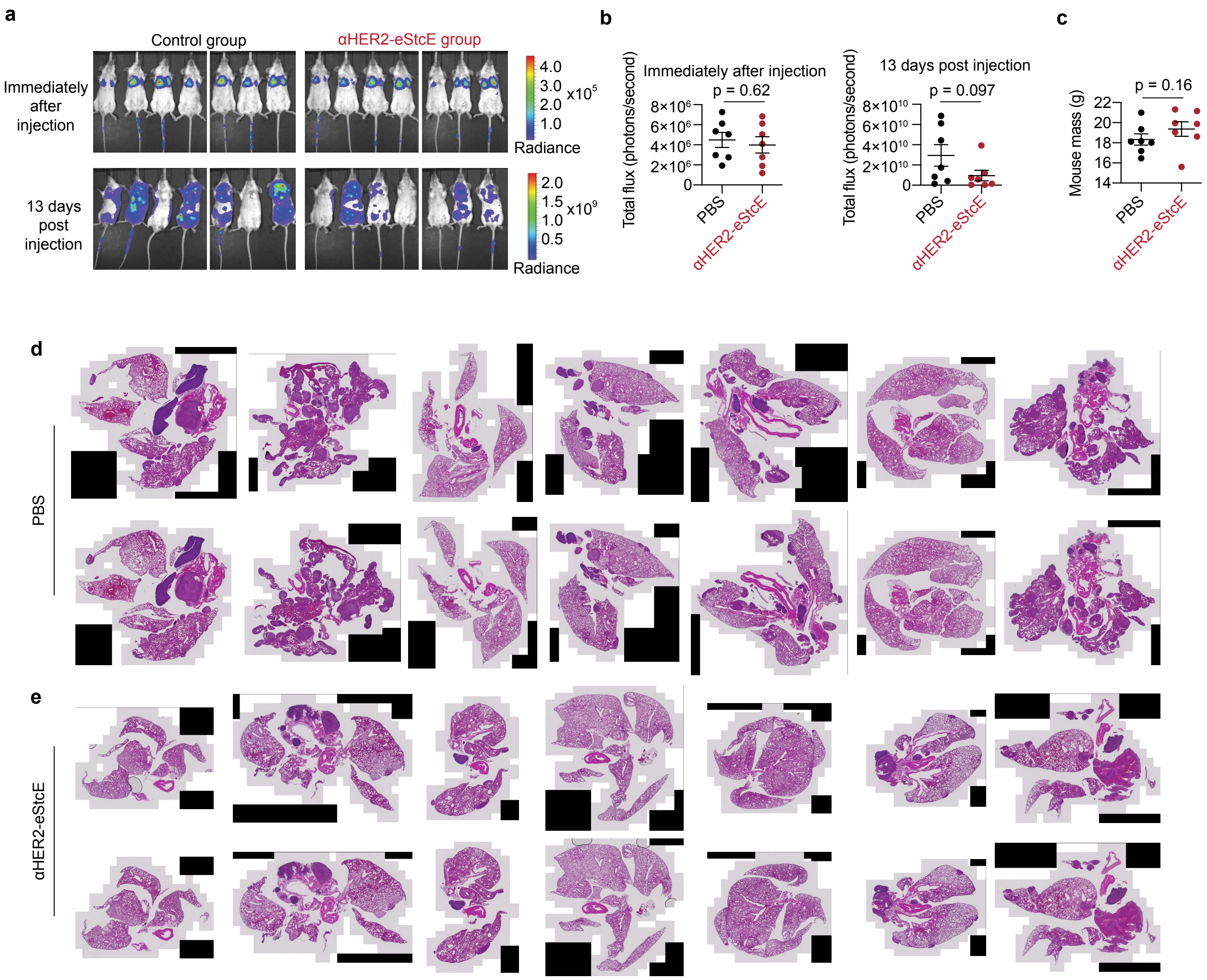


**Fig. S9. In the 4T07$^{MUC1, HER2}$ murine model of breast cancer progression, *αHER2*-eStcE reduces lung metastatic burden.**

**a,** Bioluminescent imaging of animals described in Fig. 4e-g.

**b,** Total flux measurements quantified from (**a**).

**c,** Plot depicting mouse masses of animals described in Fig. 4e-g.

**d-e,** H&E staining of lungs from PBS treated (**d**) or *αHER2*-eStcE treated (**e**) animals (*n*=7 animals per group, 2 slides per animal). Percent area of lung metastases is quantified in Fig. 4g. Data are mean ± s.e.m. *P*-values were determined using two-tailed unpaired t-test. *$p < 0.05$, **$p < 0.005$, ***$p < 0.0005$.

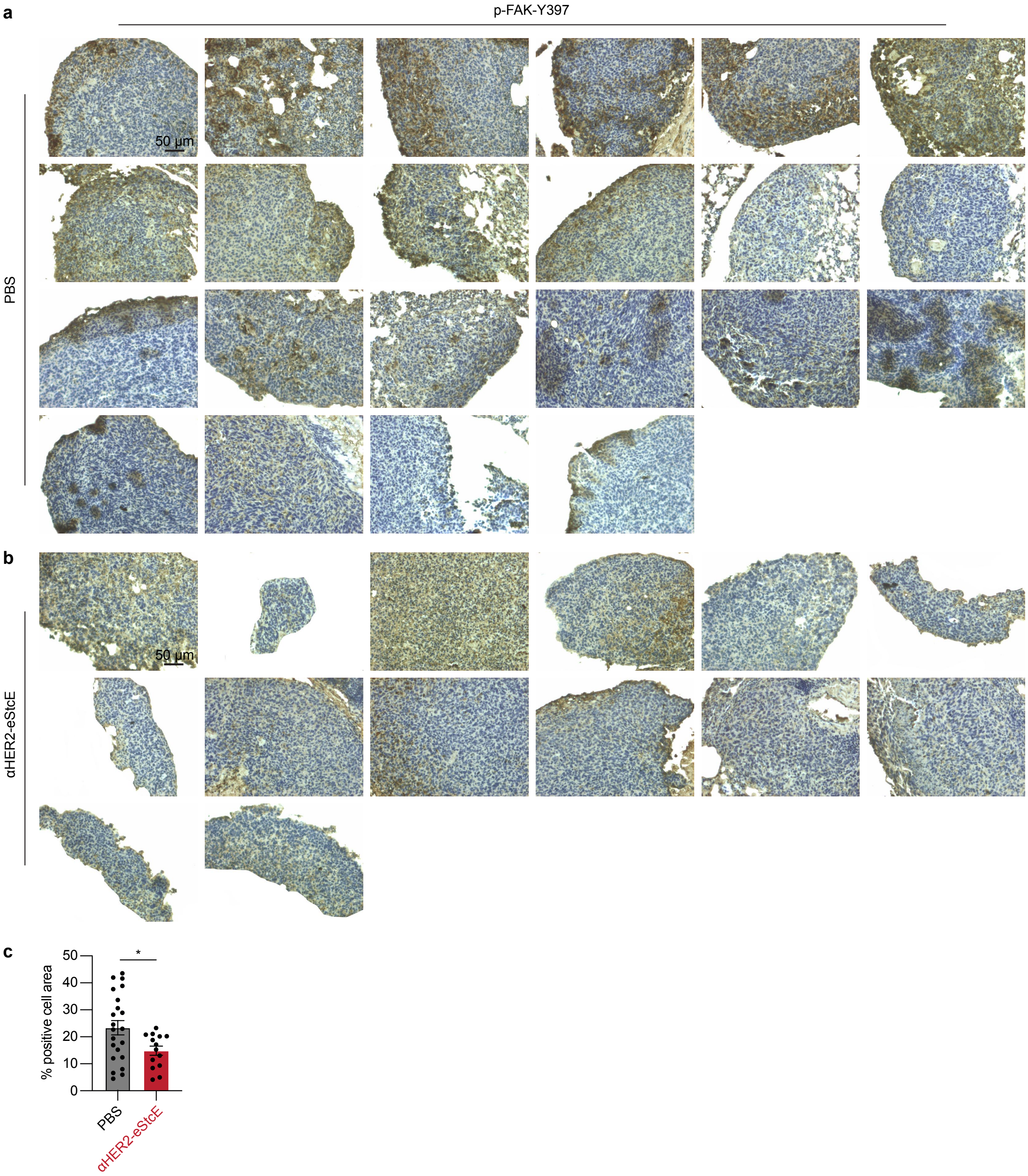


**Fig. S10. In the 4T07$^{MUC1, HER2}$ murine model of breast cancer progression, *αHER2*-eStcE reduces the prosurvival mechanosignaling marker, p-FAK-Y397.**

**a-b,** p-FAK-Y397 immunohistochemistry of lungs from PBS treated (**a**) or *αHER2*-eStcE treated (**b**) animals described in Fig. 4e-g. Each image represents a unique field-of-view.

**c,** Quantification of images from (**a-b**) using the IHC profiler plugin in ImageJ. Percent positive corresponds to positive DAB staining in the cytosol.

Data are mean ± s.e.m. *P*-values were determined using two-tailed unpaired t-test. *$p < 0.05$, **$p < 0.005$, ***$p < 0.0005$.

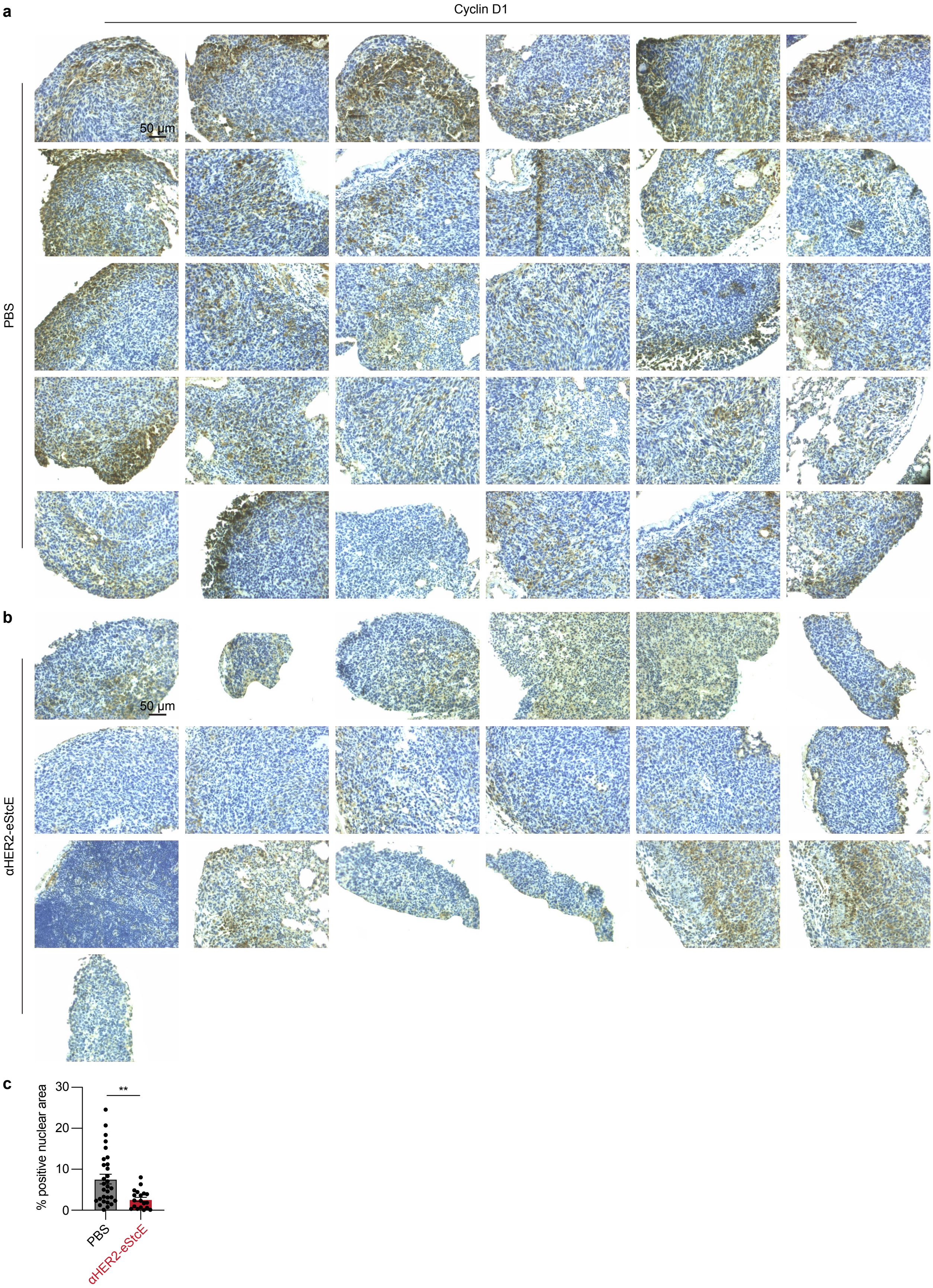
a
Cyclin D1
PBS
50 μm
b
αHER2-eStcE
50 μm
c
% positive nuclear area
30
20
10
0
**
PBS
αHER2-eStcE

**Fig. S11. In the 4T07$^{MUC1,\ HER2}$ murine model of breast cancer progression, *αHER2*-eStcE reduces the prosurvival mechanosignaling marker, cyclin D1.**
**a-b,** Cyclin D1 immunohistochemistry of lungs from PBS treated (**a**) or *αHER2*-eStcE treated (**b**) animals described in Fig. 4e-g. Each image represents a unique field-of-view.
**c,** Quantification of images from (**a-b**) using the IHC profiler plugin in ImageJ. Percent positive corresponds to positive DAB staining in the nucleus.
Data are mean ± s.e.m. *P*-values were determined using two-tailed unpaired t-test. *$p < 0.05$, **$p < 0.005$, ***$p < 0.0005$.

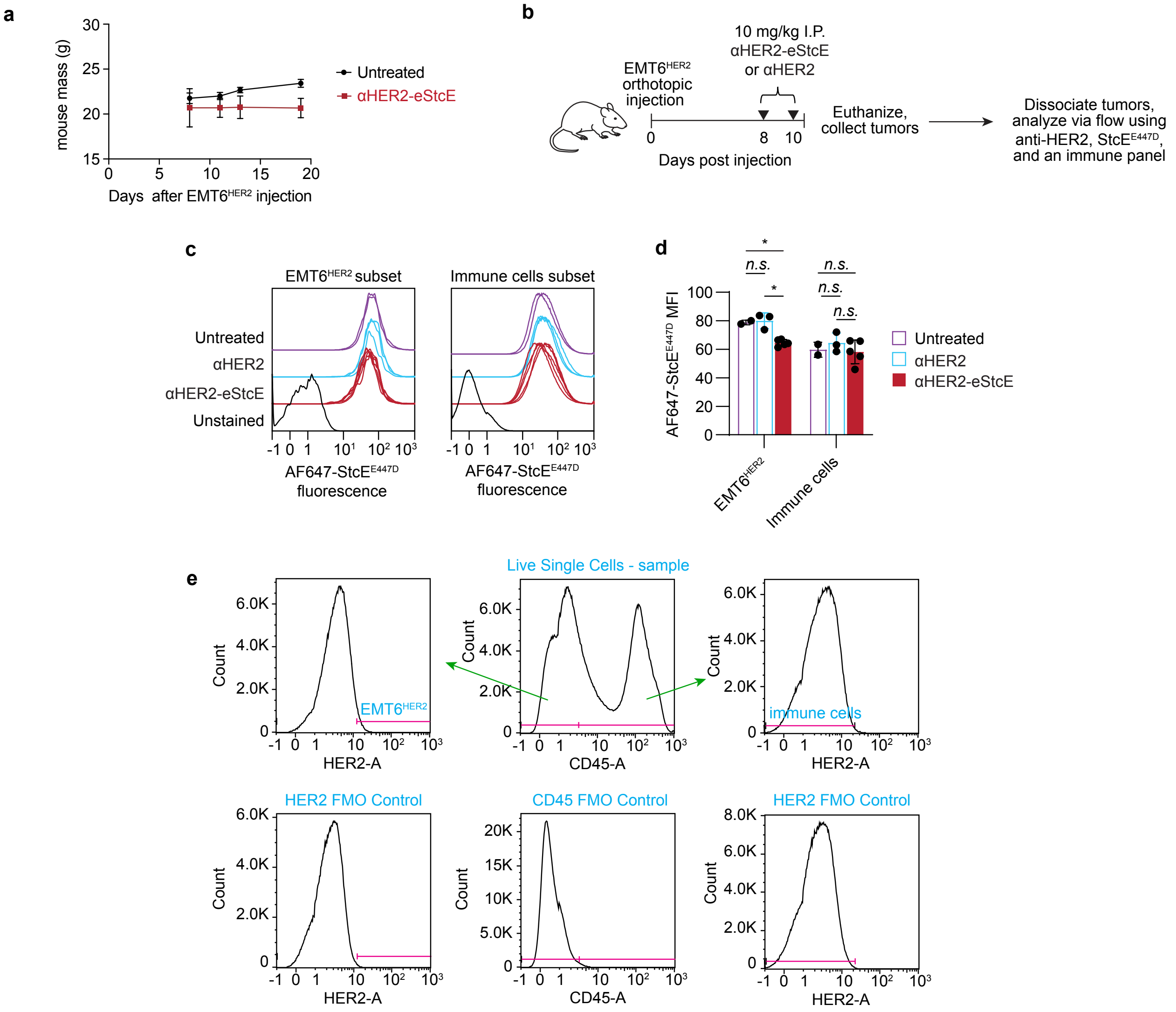


**Fig. S12. In the EMT6$^{HER2}$ murine model of breast cancer progression, *αHER2*-eStcE reduces mucin levels on EMT6$^{HER2}$ cells but not immune cells.**

**a,** Plot depicting mouse masses of animals described in Fig. 4h-j. Mouse masses for αHER2 treated mice were not measured.

**b,** Short-term treatment regimen for BALB/c mice injected with EMT6$^{HER2}$ orthotopically into the mammary fat pad. αHER2-eStcE or αHER2 were injected twice intraperitoneally (I.P.) every other day starting on day 8 ($n$=6-9 animals per group). The dose was 10 mg/kg for αHER2-eStcE and an equimolar quantity (2.8 nmol) of αHER2.

**c,** Total cell surface mucin staining of CD45$^{-}$/HER2$^{+}$ (EMT6$^{HER2}$ subset) and CD45$^{+}$/HER2$^{-}$ (immune subset) cells isolated from animals described in (**b**) using Alexa Fluor 647-labeled StcE$^{E447D}$ (AF647-StcE$^{E447D}$)[16].

**d,** Mean fluorescence intensity (MFI) values derived from (**c**) ($n$=2-5 biological replicates).

**e,** Gating strategy for (**c-d**). Fluorescence minus one controls (FMO) were used to define negative staining gates.

Data are mean ± s.e.m. (**a**) or mean ± s.d. (**d**). $P$-values were determined using Tukey-corrected two-way ANOVA. *$p < 0.05$, **$p < 0.005$, ***$p < 0.0005$.

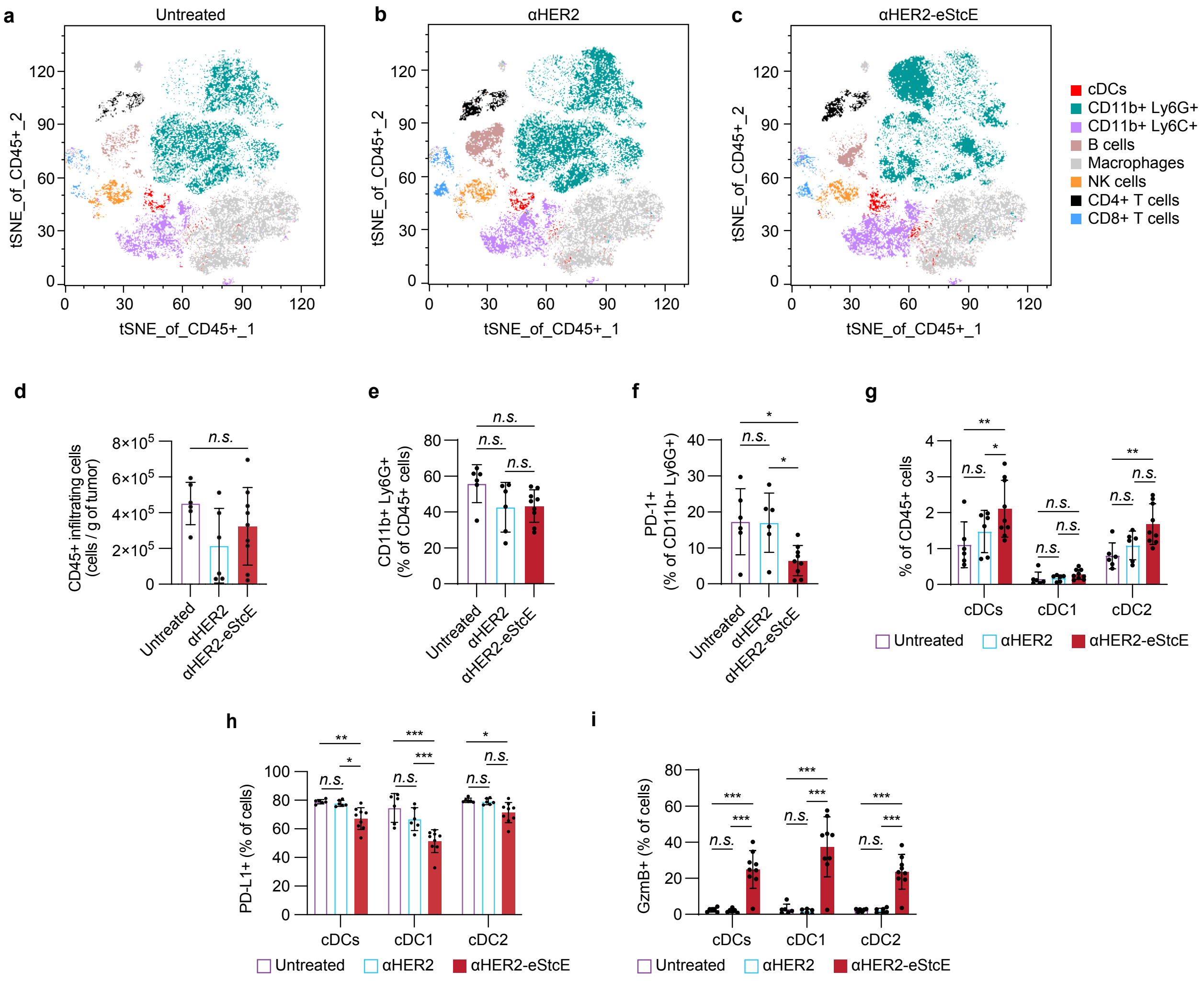


**Fig. S13. In the EMT6$^{HER2}$ murine model of breast cancer progression, *αHER2*-eStcE alters the tumor immune microenvironment.**

**a-c,** T-statistical stochastic neighbor embedding (tSNE) plots depicting immune cell subsets of tumor-infiltrating lymphocytes from untreated (**a**), αHER2 treated (**b**), and αHER2-eStcE treated (**c**) animals described in Fig. S12b. Immune subsets were defined as shown in Fig. S14.

**d,** Single, live CD45+ cells per gram of tumor.

**e,** Percent of tumor-infiltrating Ly6G+ cells as a fraction of total CD45+ cells.

**f,** Percent of PD-1+ cells in the Ly6G+ cell population.

**g,** Percent of tumor-infiltrating cDCs, cDC Type 1 (cDC1s), and cDC Type 2 (cDC2s) as a fraction of total CD45+ cells.

**h-i,** Percent of PD-L1+ (**h**) and GzmB+ (**i**) cells in cDC, cDC1, and cDC2 populations.

*n*=6-9 animals per group. Data are mean ± s.d. *P*-values were determined using Tukey-corrected one-way and two-way ANOVA. $^{*}p < 0.05$, $^{**}p < 0.005$, $^{***}p < 0.0005$.

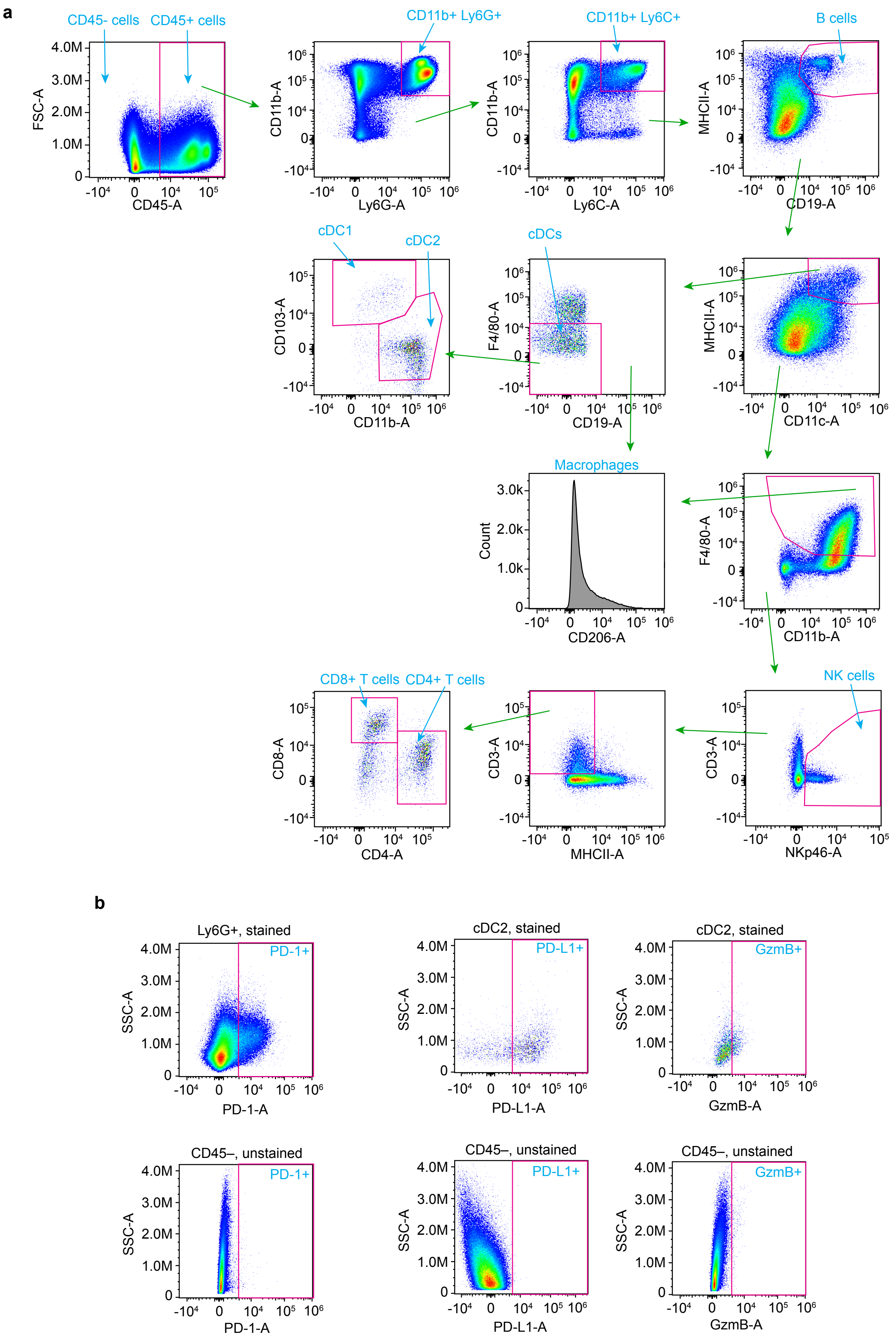
a
CD45- cells
CD45+ cells
FSC-A
CD45-A
CD11b+ Ly6G+
CD11b-A
Ly6G-A
CD11b+ Ly6C+
Ly6C-A
B cells
MHCII-A
CD19-A
cDC1
cDC2
CD103-A
CD11b-A
cDCs
F4/80-A
CD19-A
MHCII-A
CD11c-A
Macrophages
Count
CD206-A
F4/80-A
CD11b-A
CD8+ T cells
CD4+ T cells
CD8-A
CD4-A
CD3-A
MHCII-A
NK cells
CD3-A
NKp46-A
b
Ly6G+, stained
PD-1+
SSC-A
PD-1-A
cDC2, stained
PD-L1+
PD-L1-A
cDC2, stained
GzmB+
GzmB-A
CD45–, unstained
PD-1+
PD-1-A
CD45–, unstained
PD-L1+
PD-L1-A
CD45–, unstained
GzmB+
GzmB-A

**Fig. S14. Gating strategy for EMT6$^{HER2}$ immune subset profiling.**
**a,** Gating strategy used to define different immune subsets in Fig. S13 from single live cells. Plots are from a representative αHER2-eStcE treated mouse.
**b,** Defining gates for positive PD-1, PD-L1, and GzmB staining in Fig. S13f, h-i, with immune subsets from the same mouse as in (**a**). Top plots show gates on stained populations and bottom plots show gates on a fully unstained sample.
